# Dynamic multifactor hubs interact transiently with sites of active transcription in *Drosophila* embryos

**DOI:** 10.1101/377812

**Authors:** Mustafa Mir, Michael R. Stadler, Stephan A. Ortiz, Melissa M. Harrison, Xavier Darzacq, Michael B. Eisen

## Abstract

The regulation of transcription requires the coordination of numerous activities on DNA, yet it remains poorly understood how transcription factors facilitate these multiple functions. Here we use lattice light-sheet microscopy to integrate single-molecule and high-speed 4D imaging in developing *Drosophila* embryos to study the nuclear organization and interactions of the key patterning factors Zelda and Bicoid. In contrast to previous studies suggesting stable, cooperative binding, we show that both factors interact with DNA with surprisingly high off-rates. We find that both factors form dynamic subnuclear hubs, and that Bicoid binding is enriched within Zelda hubs. Remarkably, these hubs are both short lived and interact only transiently with sites of active Bicoid dependent transcription. Based on our observations we hypothesize that, beyond simply forming bridges between DNA and the transcription machinery, transcription factors can organize other proteins into hubs that transiently drive multiple activities at their gene targets.

## Introduction

The earliest stages of animal development are dominated by DNA replication and cell or nuclear division, and are primarily driven by maternally deposited RNAs and proteins. Later, control is transferred to the embryonic genome in the maternal-to-zygotic transition (MZT), during which transcription of the embryonic genome commences while maternal products are degraded (Harrison and Eisen 2015).

The MZT in *Drosophila melanogaster* begins in the early syncytial blastoderm after nine rounds of nuclear division (nuclear cycle 9, nc9) (Foe and Alberts 1983). The number of transcribed genes increases gradually as interphase periods steadily lengthen between cycles 9 and 13, before giving way to full-scale zygotic genome activation (ZGA) coincident with cellularization during the one hour long interphase of the 14th nuclear cycle (Edgar, Kiehle, and Schubiger 1986; Edgar and Schubiger 1986; Pritchard and Schubiger 1996; Anderson and Lengyel 1981; Zalokar 1976).

Thousands of genes become transcriptionally active during the MZT, including several hundred transcribed in defined spatial and temporal patterns along the anterior-posterior (AP) and dorsal-ventral (DV) axes (Combs and Eisen 2017, 2013; Lécuyer et al. 2007; Wilk et al. 2016; Tomancak et al. 2007), which serve as the first markers of the nascent body plan of the developing embryo. The formation of these patterns is directed through interactions of DNA-binding proteins known as transcription factors with non-coding regulatory genomic regions known as enhancers. Enhancers are typically bound by combinations of activating and repressing transcription factors and drive transcription of target genes in patterns that depend on the differential combination of factors present in nuclei at different positions within the embryo. However, beyond this basic paradigm, it remains poorly understood how the composition and arrangement of transcription factor binding at enhancers dictates the output of the genes they regulate and what role interactions among binding factors play in this process.

In recent years, it has become clear that patterning transcription factors are only part of the complex systems that specify enhancer activity. Among the key additional players is the ubiquitously distributed maternal factor Zelda (Staudt 2006; ten Bosch, Benavides, and Cline 2006; Liang et al. 2008; De Renzis et al. 2007) (Zld, also known as Vielfaltig, Vfl) that we and others have shown plays a central role in the spatiotemporal coordination of gene activation, and in facilitating the binding of patterning factors to their target enhancers (Harrison et al. 2011; Li et al. 2014; Nien et al. 2011; Foo et al. 2014; Schulz et al. 2015; Sun et al. 2015).

Zelda is often described as a “pioneer” transcription factor (Zaret and Mango 2016), in that its primary function appears to be to facilitate the binding of other factors indirectly by influencing chromatin state. However, how it accomplishes this remains unclear. Zelda has a cluster of four C2H2 Zn-fingers near the C-terminus that mediate its DNA binding activity and 2 additional ZFs near the Nterminus which have been implicated in controlling its activation potential (Hamm et al. 2017), but most of the rest of the protein consists of varying types of low-complexity sequence.

Such low-complexity domains (LCDs) are thought to facilitate protein-protein interactions that mediate the formation of higher order structures, including phase separated domains (Kato and McKnight 2018; Brangwynne et al. 2009). There is increasing evidence that higher-order structures mediated by low-complexity domains play an important role in transcriptional regulation (Chong et al. 2018; Boehning et al. 2018; Strom et al. 2017; Kato and McKnight 2018), although the precise nature of this role remains less than clear. One hypothesis is that domains formed by homo- and heterotypic interactions between LCDs serve to locally enrich transcription factors, potentially in the vicinity of their targets (Tsai et al. 2017), thereby altering their local concentration and modulating their binding dynamics.

We recently explored this idea by utilizing lattice light-sheet microscopy (LLSM) (B.-C. Chen et al. 2014) to carry out single-molecule imaging and tracking of eGFP labeled Bicoid (Bcd)—the primary anterior morphogen in *D. melanogaster*—in living embryos (Mir et al. 2017). Bicoid proteins are distributed in a concentration gradient along the anterior-posterior axis, and activate approximately 100 genes in a concentration dependent manner, primarily in anterior portions of the embryo (Z. Xu et al. 2014). The sharpness of the responses of Bcd targets to its gradient has led to the proposal of various models of cooperative regulation (Frohnhöfer and Nüsslein-Volhard 1986; Driever and Nüsslein-Volhard 1988), but the molecular basis for this apparent cooperation remains incompletely worked out.

We previously showed that Bcd binds DNA transiently (has a high *k*_off_) and that its binding is concentrated in discrete, sub-nuclear domains of locally high Bcd density that we refer to as “hubs” (Mir et al. 2017). These hubs are more prominent in posterior nuclei where Bcd concentration is low but in which it still binds specifically to target loci. We proposed that Bcd hubs facilitate binding, especially at low concentration, by increasing the local concentration of Bcd in the presence of target loci, thereby increasing *k*_on_ and factor occupancy (Mir et al. 2017).

Based on previous observations (Hannon, Blythe, and Wieschaus 2017) that Bcd binding in more posterior nuclei is dependent on Zld, we examined the distribution of Bcd binding in nuclei lacking Zld and found that Bcd hubs no longer form (Mir et al. 2017). Our preliminary experiments with fluorescently tagged Zld revealed that it also forms hubs (distinct clusters of Zld were also recently reported by (Dufourt et al. 2018)).

The combined observations that Bcd forms hubs that depend on the presence of Zld, and that Zld also forms hubs motivated us to quantify the spatial and temporal relationships between Zld and Bcd molecules in *Drosophila* embryos. However, these experiments required several advancements in our technical capabilities to both tag and image single molecules.

Here we first describe Cas9-mediated tagging of endogenous loci with bright, photoswitchable fluorescent proteins that provide greatly improved signal-to-noise and tracking abilities, modifications to the LLSM necessary to activate these tagged proteins, and the development of biological and analytical tools to study the interactions between proteins and also between proteins and sites of active transcription. We use this technological platform to characterize the singlemolecule and bulk behavior of Zld and Bcd in isolation, in relation to each other and to the transcriptional activation of the canonical Bcd target gene, *hunchback* (*hb*).

We find that both Bcd and Zld bind DNA highly transiently, with residence times on the order of seconds. Furthermore, both proteins form high-concentration hubs in interphase nuclei which are highly dynamic and variable in nature. By simultaneously imaging the bulk spatial distribution of Zld (to track hubs) and single molecules of Bcd, we show that Bcd binding is both enriched and stabilized within Zld hubs, an effect that becomes more pronounced at low Bcd concentrations in the embryo posterior. Finally, we explore the functional role of Zld and Bcd hubs in activating *hb* and find that hubs of both proteins interact transiently with the active *hb* locus, with preferential interactions of Bcd hubs with active loci leading to a time-averaged enrichment of the protein at the locus. Collectively our data suggest a model in which dynamic multi-factor hubs regulate transcription through stochastic encounters with target genes.

## Results

### Single-molecule tracking of proteins endogenously tagged with photoactivatable fluorescent proteins

We used Cas9-mediated homologous replacement (Bassett et al. 2013, 2014; Gratz et al. 2013, 2014; Yu et al. 2013; Baena-Lopez et al. 2013; Sebo et al. 2013; Kondo and Ueda 2013; Ren et al. 2013) (Figure 1 - figure supplement 1) to tag endogenous loci of Bcd and Zld at their N-termini with the photoactivatable fluorescent protein mEos3.2 (Zhang et al. 2012), which has high-quantum efficiency, is highly monomeric, and photostable compared to other photoactivatable proteins. Zld was also independently tagged with the bright green fluorescent protein mNeonGreen (Hostettler et al. 2017). These lines are homozygous viable and have been maintained as homozygous lines for many generations. To serve as a controls for single-molecule experiments, we also generated lines containing ubiquitously expressed mEos3.2-tagged Histone H2B (His2B).

To utilize the photoactivatable mEos3.2 for single-molecule tracking, we modified a lattice light-sheet microscope (B.-C. Chen et al. 2014) to allow continuous and tunable photoactivation from 405 nm laser (Figure 1 - figure supplement 2; Video 1). We optimized this setup using mEos3.2-Zld, controlling particle density (Figure 1 - figure Supplement 3) to facilitate tracking (Hansen et al. 2018; Izeddin et al. 2014), and found that we could obtain excellent signal to noise ratios sufficient for robust single-molecule detection (Figure 1 A and Videos 2-4) and tracking of both mobile and immobile molecules at frame intervals ranging from 10 to 500 ms (Figure 1B and Videos 56).

**Figure 1.**
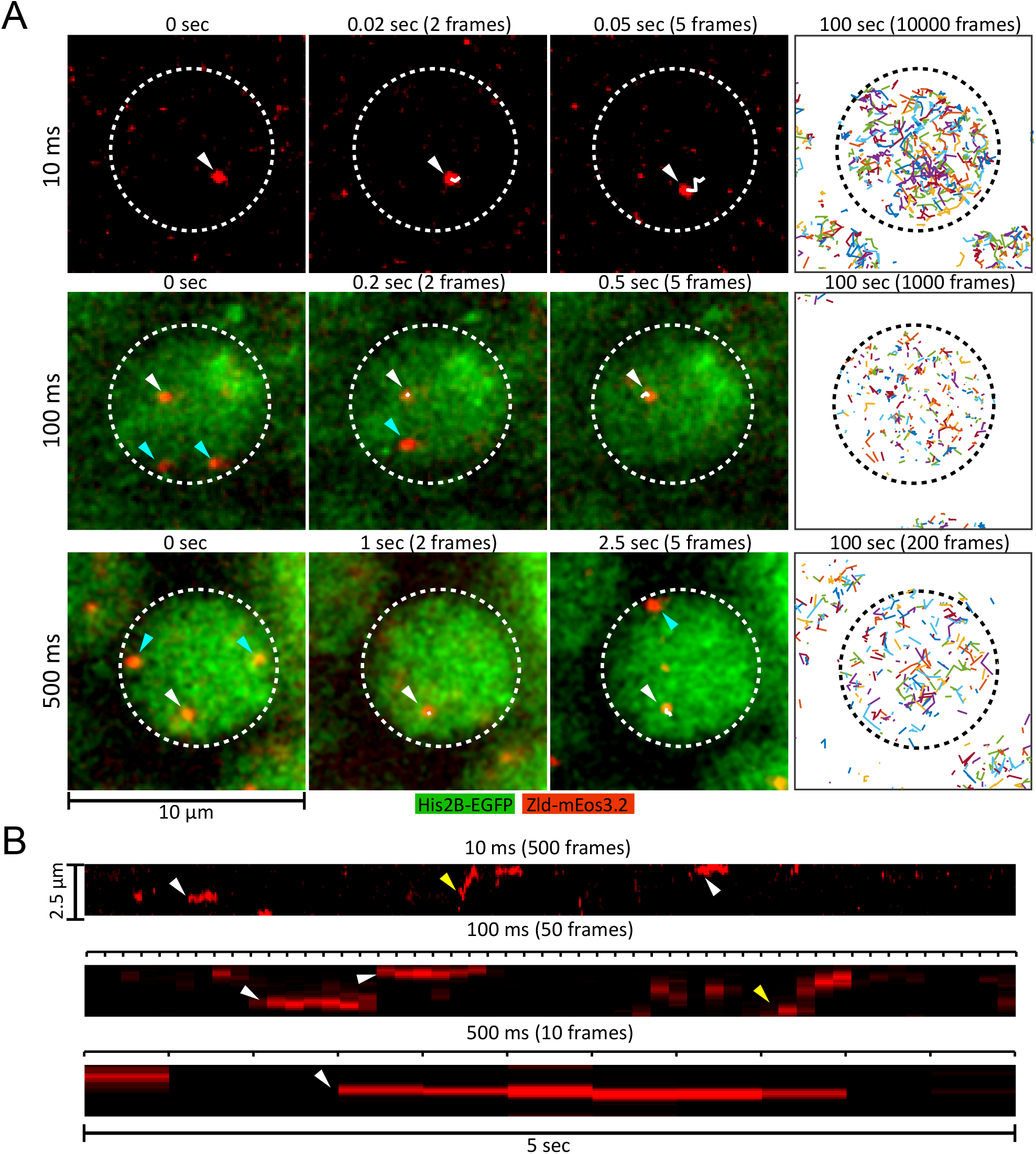
Live embryo single molecule imaging and tracking of endogenous mEos3.2-Zld. **(A)** First 3 columns are example images showing single molecules of mEos3.2-Zld tracked over at least 5 frames (white arrows and trajectories) at frame rates of 10, 100 and 500 ms. Cyan arrows indicate molecules that appear for only 1 frame and are thus detected but not tracked. For the 100 and 500 ms data enough signal is present in the His2B-eGFP channel from the activation laser to enable simultaneous imaging of chromatin. Last column shows all single molecule trajectories acquired in each nucleus over 100 sec, corresponding to 539, 263, and 186 trajectories over 10000, 1000, and 200 frames for the 10, 100 and 500 ms data respectively. Dotted lines indicate the boundary of a nucleus. Contrast was manually adjusted for visualization. **(B)** Representative kymographs over 5 seconds of imaging, corresponding to 500, 50, and 10 frames for the 10, 100 and 500 ms frame rate data respectively. Yellow arrows point to molecules that display relatively large motions, and white arrows to immobile molecules.

**Figure 1 - figure supplement 1.**
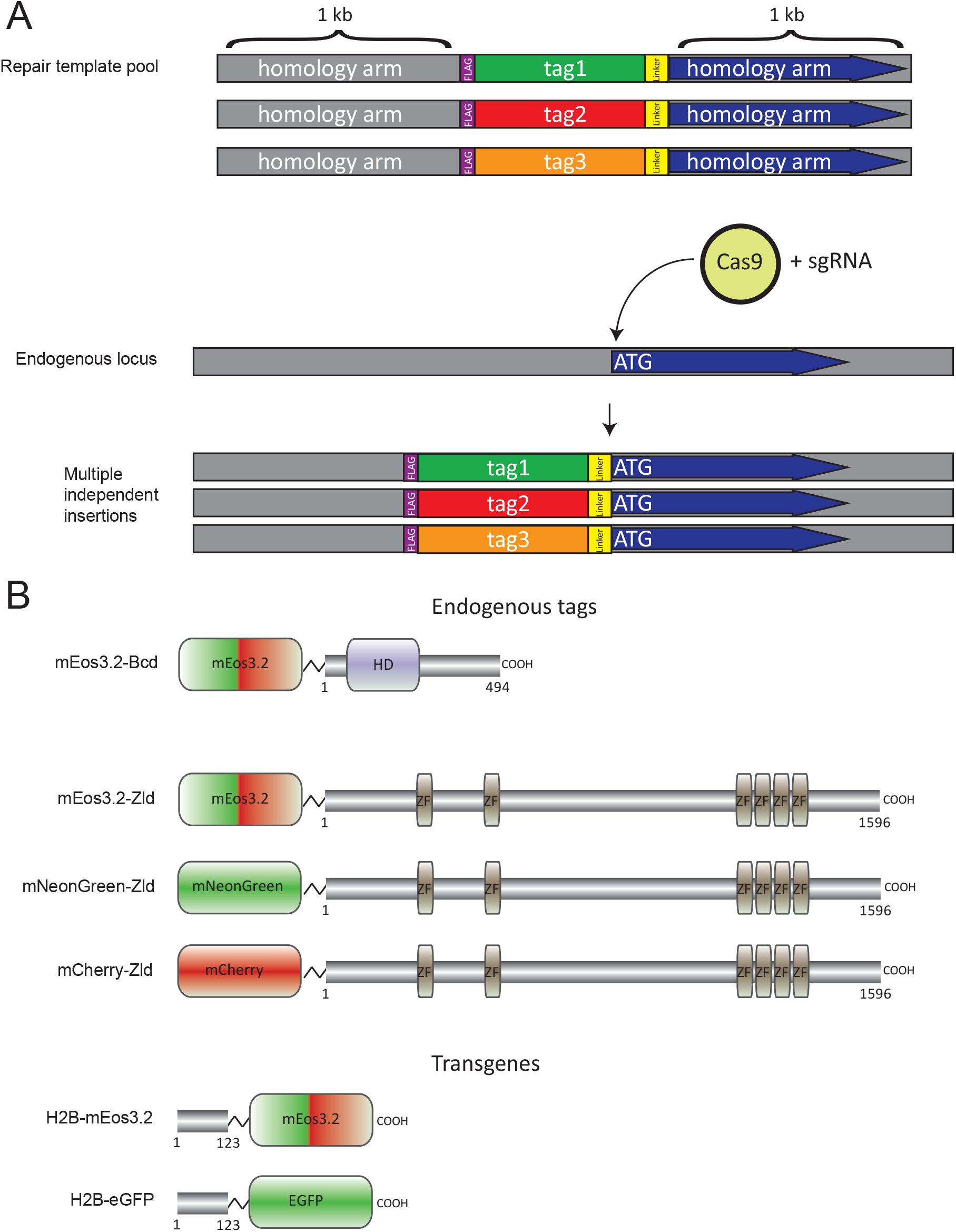
Overview of CRISPR-Cas9 genome editing strategy. **(A)** Pools of homology repair template plasmids containing different protein tags were co-injected with a plasmid encoding an sgRNA targeting the N-terminus of the target coding sequence into embryos expressing transgenic Cas9. PCR genotyping and DNA sequencing were used to find chromosomes containing tag insertions and to determine the identity of the inserted protein. **(B)** Fusion proteins used in this study. Zelda and Bicoid fusion proteins were generated by inserting fluorescent proteins at the endogenous locus. To avoid potential problems associated with endogenous tagging of a multicopy locus, histone fusions were supplied as exogenous transgenes.

**Figure 1 - figure supplement 2.**
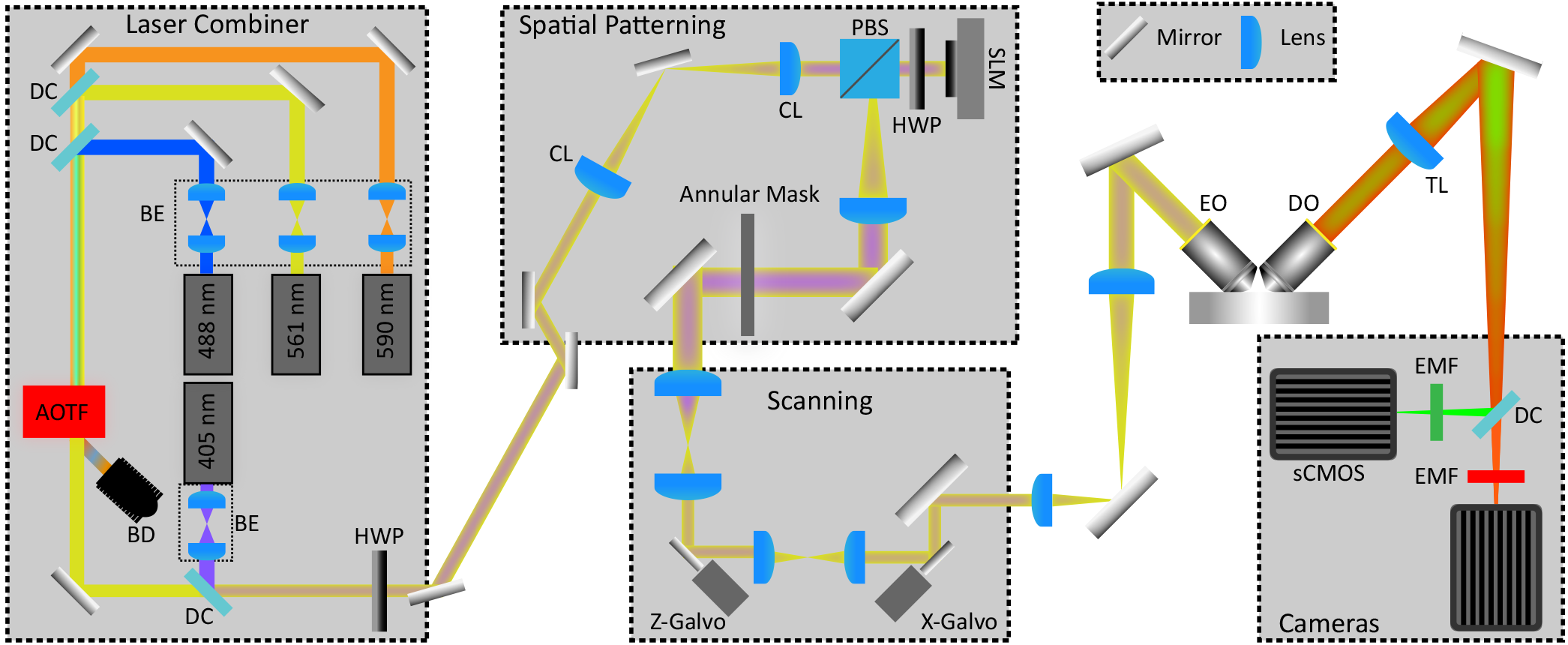
Simplified Schematic of Lattice Light Sheet Microscope. The schematic is organized to show the major modules of the microscope. The Laser Combiner module contains 6 lasers (3 shown here) for excitation ranging from 405 to 639 nm, each of which are independently expanded and collimated by using a pair of lenses that serve as a beam expander (BE). The paths of each laser are combined and made collinear by using 1 mirror and 1 dichroic mirror (DC) per laser. The combined beams are then input into an Acousto-optic tunable filter (AOTF) for rapid switching between lasers and control of power. A beam dump (BD) is used to safely capture the light from lasers not being used. To achieve constant and controllable photoactivation the laser combiner was modified to include a 405 nm laser that bypasses the AOTF. A half-wave plate (HWP) is used to adjust the polarization of the input light to the Spatial Patterning module. For spatial patterning, a pair of cylindrical lenses (CL) are used to stretch the Gaussian beam output from the Laser Combiner module into a thin stripe, which illuminates the Spatial Light Modulator (SLM) after passing through a polarizing beam splitter (PBS) and a second HWP. A lens projects the Fourier transform of the plane of the SLM onto an annular mask which is used to confine the spatial frequencies of the patterned light to the desired minimum and maximum numerical apertures. In the scanning module, a pair of lenses de-magnifies and projects the annular mask plane onto first the z-galvo scanning mirror for moving the light sheet through the sample, and a second pair-of lenses relays the plane of the z-galvo onto the x-galvo for dithering the sheet for uniform illumination. Another pair of lenses is then used to project a magnified image of the galvo planes to the back focal plane of the excitation objective (EO) which focuses the light to project the lattice pattern through the sample. An orthogonally placed detection objective (DO) collects the emission light, and a tube-lens (TL) then forms an image at each cameras’ sensor plane. A dichroic mirror first splits the light into red (>560 nm) and green (<560 nm) channels, followed by a narrower bandpass emission filter (EMF) for further filtering before each camera. With the exception of the modifications to the laser combiner module and the use of two Hamamatsu sCMOS ORCA Flash 4.0 cameras for detection the design is identical to what was originally described by (Chen et al., 2014).

**Figure 1 - figure supplement 3.**
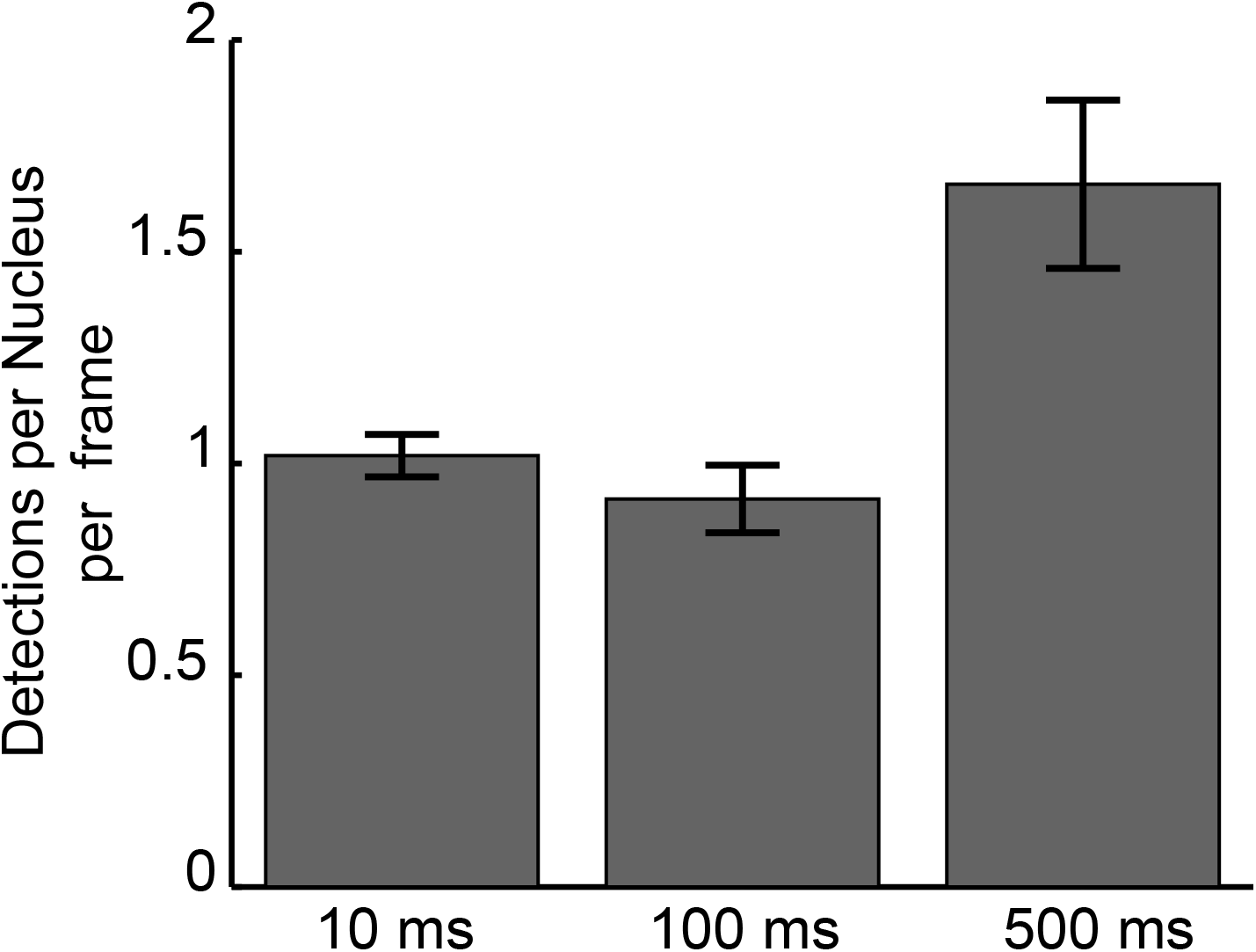
Mean detections per nucleus per frame for each frame rate. Detections per nucleus/frame characterized over 804245, 78281, 15165 frames of imaging and 169, 434, and 359 nuclei for 10 ms, 100 ms, and 500 ms, datasets respectively. Error bars show standard errors over all nuclei.

We deployed this platform to perform single molecule imaging and tracking of Zld, Bcd, and His2B at 10, 100 ms, and 500 ms frame intervals (Figure 2, Videos 7-9). These different temporal resolutions each capture distinct aspects of molecular behavior: short exposure times are sufficient to detect single molecules, but fast enough to track even rapidly diffusing molecules (Video 7). However, because imaging single molecules at high-temporal resolution (10 ms) requires high-excitation illumination, most bound molecules photobleach before they unbind, encumbering the accurate measurement of long binding times. At longer exposure times of 100 ms and 500 ms, fast diffusing proteins are blurred into the background, and lower illumination lowers photobleaching rates such that unbinding events can be detected (Hansen et al. 2017; Mir et al. 2017; Normanno et al. 2015; J. Chen et al. 2014; Mazza et al. 2012). Thus we use 10 ms data to measure the diffusion characteristics of bound and unbound molecules, as well as to determine the fraction of total molecules that are bound (Hansen et al. 2018), and 100 and 500 ms data to measure the duration and spatial distribution of binding events.

To gain an understanding of the dynamics of a protein which is stably associated with chromatin, we first examined single molecule trajectories of the histone His2B at all three temporal scales. Histones are widely used as a benchmark for stably bound molecules (Mazza et al. 2012; Hansen et al. 2017; Teves et al. 2016), and we validate that His2B is a suitable control in the early *Drosophila* embryo through fluorescence recovery after photobleaching measurements (FRAP) (Figure 2-figure supplement 1). Consistent with this stable association, a visual examination of the single particle trajectories of His2B at 10 ms frame rates illustrate that the vast majority of His2B molecules are immobile and confined within the localization accuracy of our measurements (Figure 2, top left and Video 7). In comparison, the Zld and Bcd trajectories at 10 ms frame rates exhibit motions consistent with a mixed population of both chromatin bound and mobile molecules (Figure 2 left column and Video7).

**Figure 2.**
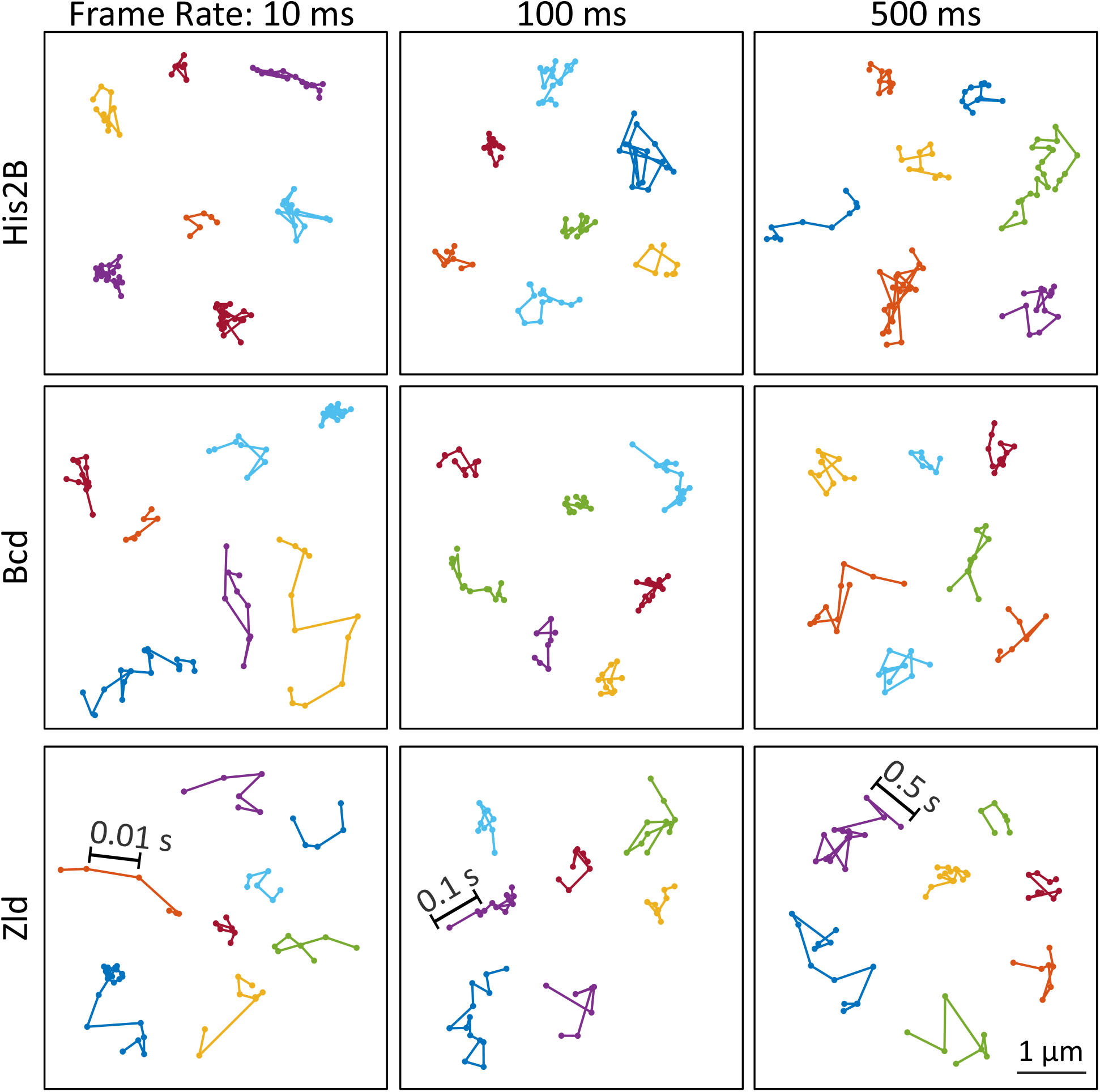
Representative single molecule trajectories of His2B, Bcd and Zelda. Representative single molecule trajectories of His2B, Bcd, and Zld from data acquired at frame rates of 10, 100 and 500 ms.

**Figure 2-figure supplement 1.**
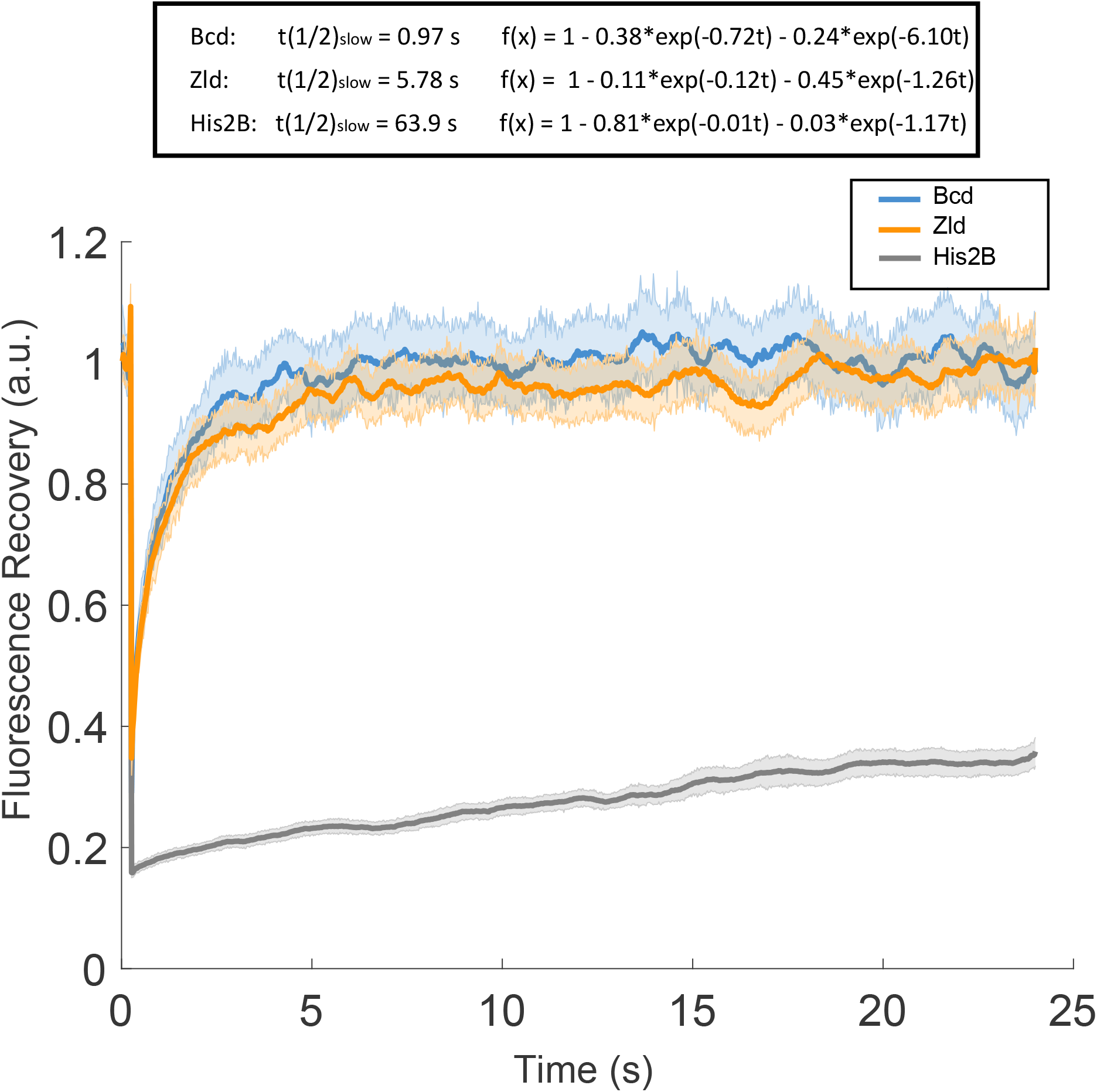
Fluorescence Recovery After Photobleaching. Fluorescence Recovery After Photobleaching (FRAP) curves for Bcd, Zld, and His2B and results of 2 exponential fitting. Solid lines are averages over at least 27 measurements and shaded regions indicate standard error.

When tracked over several seconds using exposure times of 10 and 500 ms (Figure 2, middle and right), the His2B trajectories now reflect the underlying motion of chromatin. We note a significantly greater apparent chromatin motion in early *Drosophila* embryos than is observed in mammalian cells in interphase where histones typically exhibit mobility less than the achievable localization accuracy (Hansen et al. 2018). At these slower frame rates molecules of Zld and Bcd which are not immobile for a significant portion of the exposure time motion blur into the background (Watanabe and Mitchison 2002; Hansen et al. 2017; Zhang et al. 2012). As a result, the trajectories of all 3 proteins now appear visually similar with the exception that His2B trajectories are longer in time due to their stable interaction with chromatin (Video 9), with the length of trajectories now limited by unbinding, defocalization, and photobleaching. Having established His2B as a suitable control for a largely chromatin-bound protein we next quantify and compare the single molecule dynamics of Zld and Bcd in order to gain insight on how they explore the nucleoplasm and bind to DNA to regulate transcription.

### Bicoid and Zelda bind transiently and have large free populations

We first quantified the fraction of molecules bound, and the diffusion coefficients of free and bound molecules for His2B, Zld and Bcd, by analyzing the distributions of displacements (Hansen et al. 2018) from the high speed (10 ms frame rate) data (Figure 3A and Figure 3 - figure supplement 1). Visually the displacement distributions indicate that a greater fraction of both Zld and Bcd molecules are mobile (Figure 3A, displacements > 150-200 nm) than for His2B.

To quantify the single-molecule kinetics of all 3 proteins, the displacement distributions were fit to a 2-state (diffusing or bound) kinetic model (Figure 3 - figure supplement 1) assuming Brownian motion under steady-state conditions and taking into account effects from localization errors and defocalization bias (Mazza et al. 2012; Hansen et al. 2017, 2018). We find that ~50 % of Zld and Bcd, and 88 % of His2B molecules are bound (Figure 3B and Figure 3 - figure supplement 1). The mean bound diffusion coefficient for His2B is lowest followed by Bcd, and Zld, whereas the free diffusion coefficients for Zld are slightly lower than both Bcd, and His2B (Figure 3 - figure supplement 1). The ~50% bound population of Zld and Bcd indicate that both proteins spend roughly the same amount of time on nuclear exploration (searching for a binding target) and actually binding to chromatin (Hansen et al. 2017).

Next we calculated the survival probability (the probability of trajectories lasting a certain amount of time) for the three factors at all three frame rates (Figure 4A). At all frame intervals, the length of His2B trajectories are, on average, longer than those of Zld and Bcd (Figure 4A). These longer trajectories reflect the greater fraction of bound His2B molecules as they defocalize with a lower probability. Since we expect, on average, the effects of nuclear and chromatin motion, as well as photobleaching, to be consistent for data acquired on the bound population of all three proteins, the longer His2B trajectories show both that His2B binds for longer than Bcd or Zld, and that unbinding and not photobleaching is likely to be dominant for Bcd and Zld trajectories at 500 ms exposure times, allowing us to estimate residence times.

**Figure 3.**
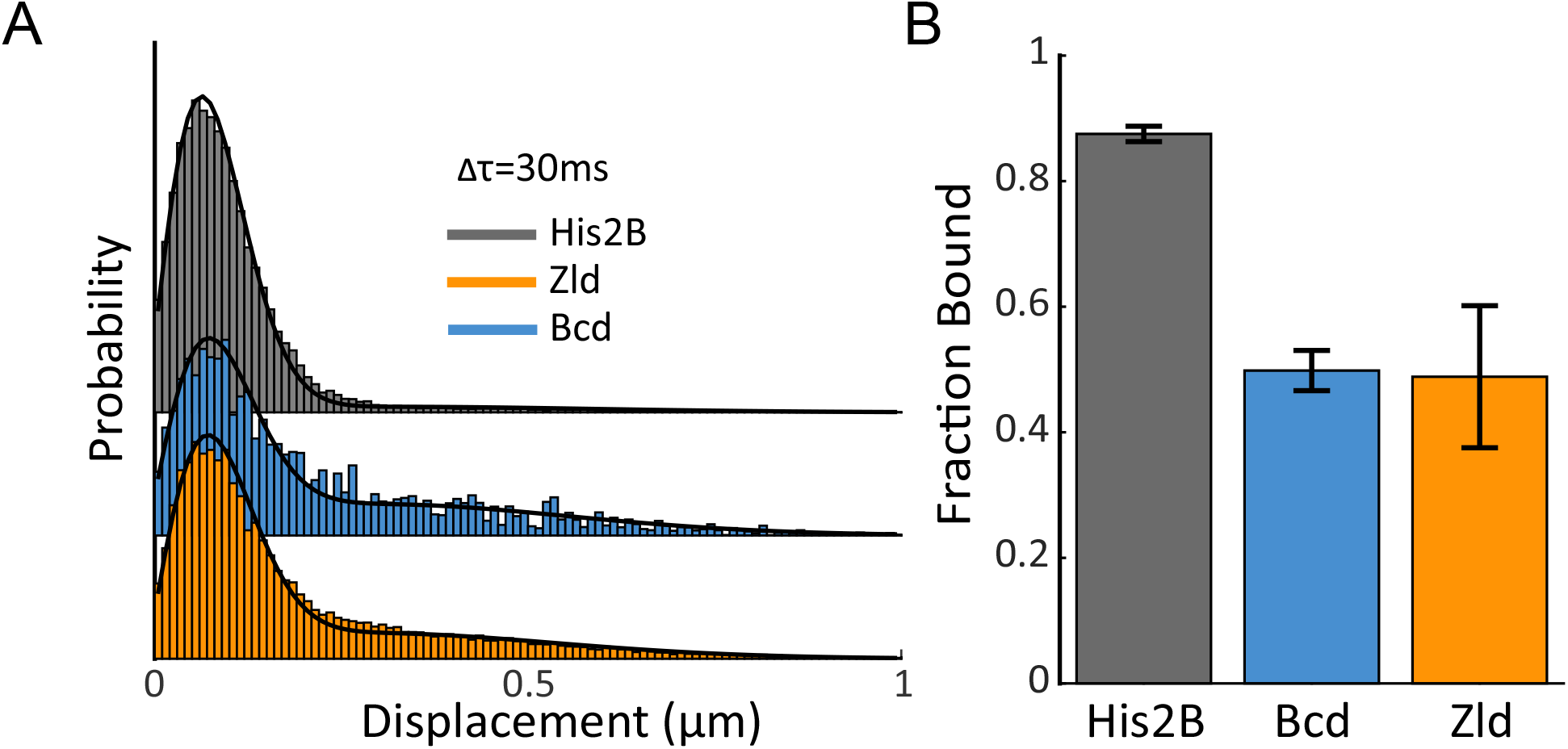
Fraction bound of His2B, Zld, and Bcd molecules. **(A)** Histograms of displacements for His2B, Zld, and Bcd after 3 consecutive frames (Δ_T_=30ms) at a frame rate of 10 ms. The Zld and Bcd distributions show a right tail indicative of a large free population that is missing from His2B distribution. Black lines are fits from 2-state kinetic modelling, data shown is compiled from 3 embryos totalling 77869, 81660, and 11003 trajectories and 30, 128, 41 nuclei for His2B, Zld, and Bcd respectively. **(B)** Fraction of molecules bound as determined from kinetic modelling of the displacement distributions, a summary of the model parameters is shown in Figure 3-figure supplement 1. Error bars are standard errors over 3 embryos.

**Figure 4.**
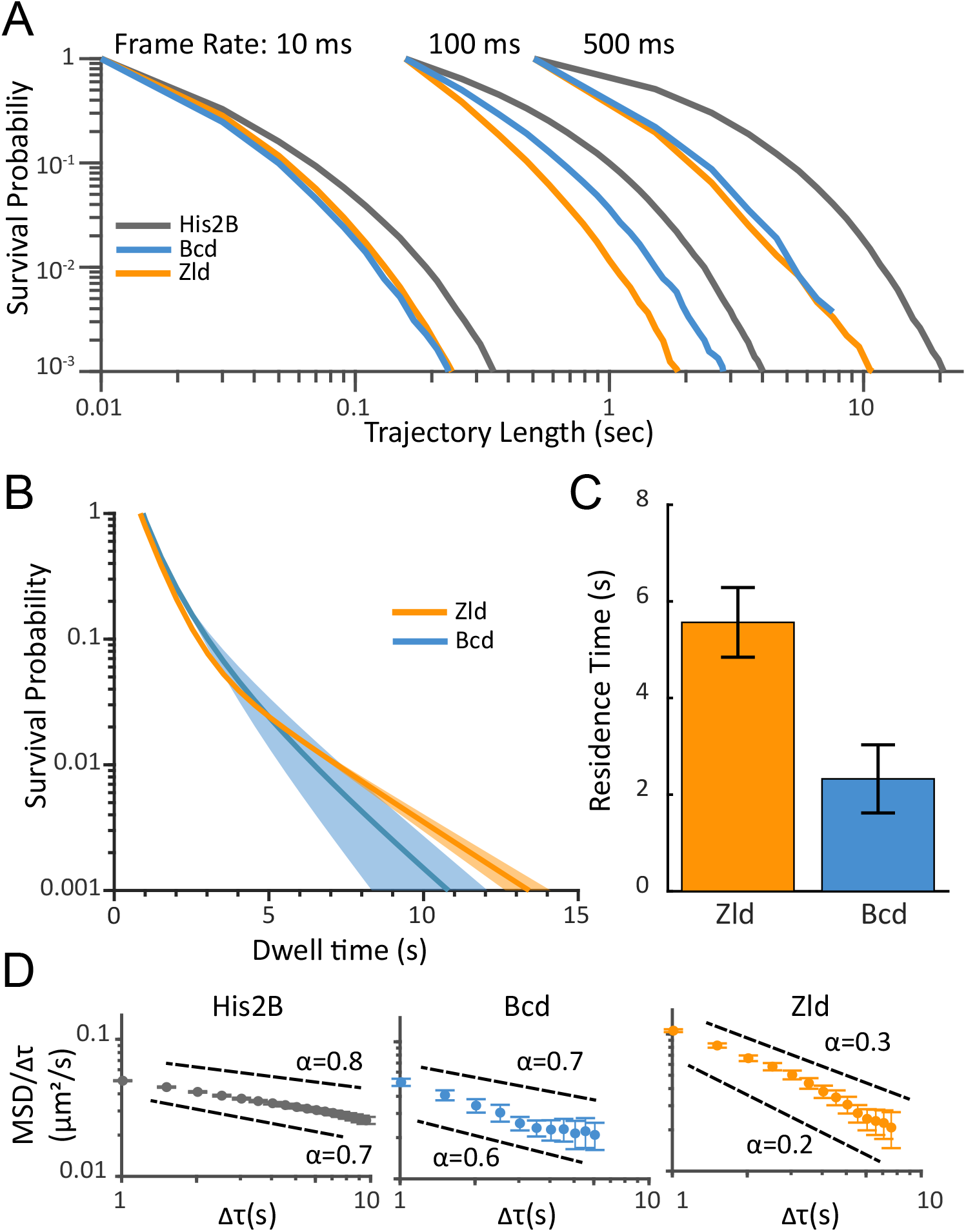
Residence times and dynamics of bound Zld and Bcd molecules. **(A)** Raw survival probabilities of trajectories as function of length at all frame rates. Calculated over 77869, 81660, 11003, at 10 ms, 107998, 42698, 8990 at 100 ms, and 2420, 14487, 47681 at 500 ms, trajectories for His2B, Zld, and Bcd respectively. **(B)** Uncorrected (for photobleaching and defocalization) two-exponent fits to the survival probability distributions obtained from the 500 ms frame rate data. Dark solid lines are the mean over fits from 3 embryos and the shaded regions indicated the standard error. **(C)** Bias corrected (for photobleaching and defocalization) quantification of the slow residence times for Zld (5.56 ± 0.72 s) and Bcd (2.33 ± 0.71 s). Error bars indicate standard error over 3 embryos for a total of 188 and 171 nuclei for Bcd and Zld respectively. **(D)** MSD/⊤ curves for His2B, Bcd, and Zld at 500 ms frame rates plotted on log-log-scale. For anomalous diffusion MSD(⊤)= Γ⊤α, where α is the anomalous diffusion coefficient. For MSD/⊤, in log-log space, the slope is thus 0 for completely free diffusion that is when α=1, and sub-diffusive(0<α<1), motions display higher negative slopes, with lower α corresponding to more anomalous motion.

**Figure 4 - figure supplement 1.**
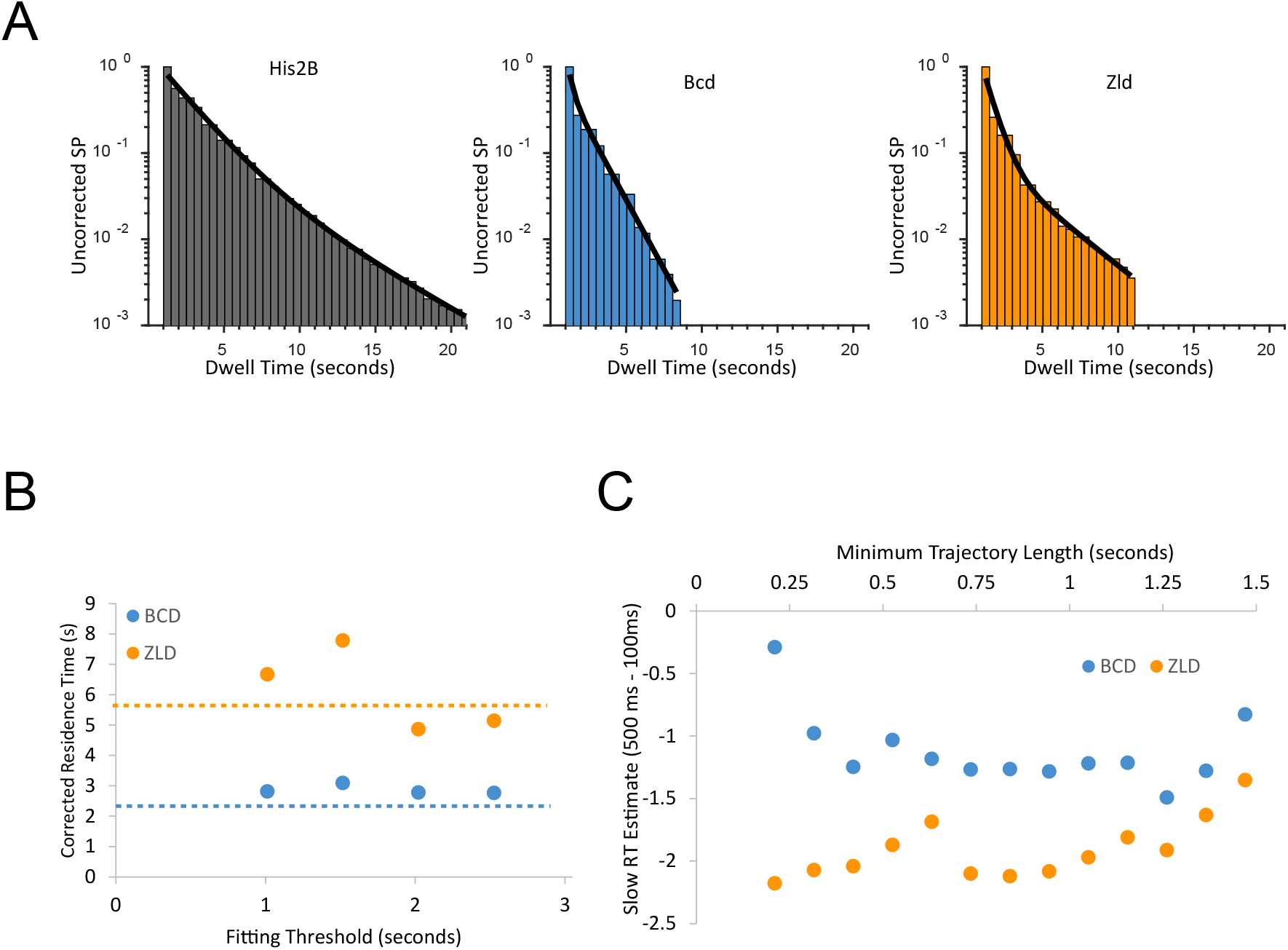
Inference of residence times from single molecule trajectories. **(A)** Example fits of two-exponent model (black line) to the surival probability (SP) distributions from a single embryo for each protein studied **(B)** Bias corrected residence time (RT) as a function of the minimum trajectory length threshold used for fitting, the reported inferred residence time is from when the fit parameter plateaus at a threshold of 2.02 s for both proteins **(C)** Comparison of inferred residence times (RT) from 100 ms vs 500 ms frame rate data sets as a function of the minimum trajectory length fitting threshold.

**Figure 4 - figure supplement 2.**
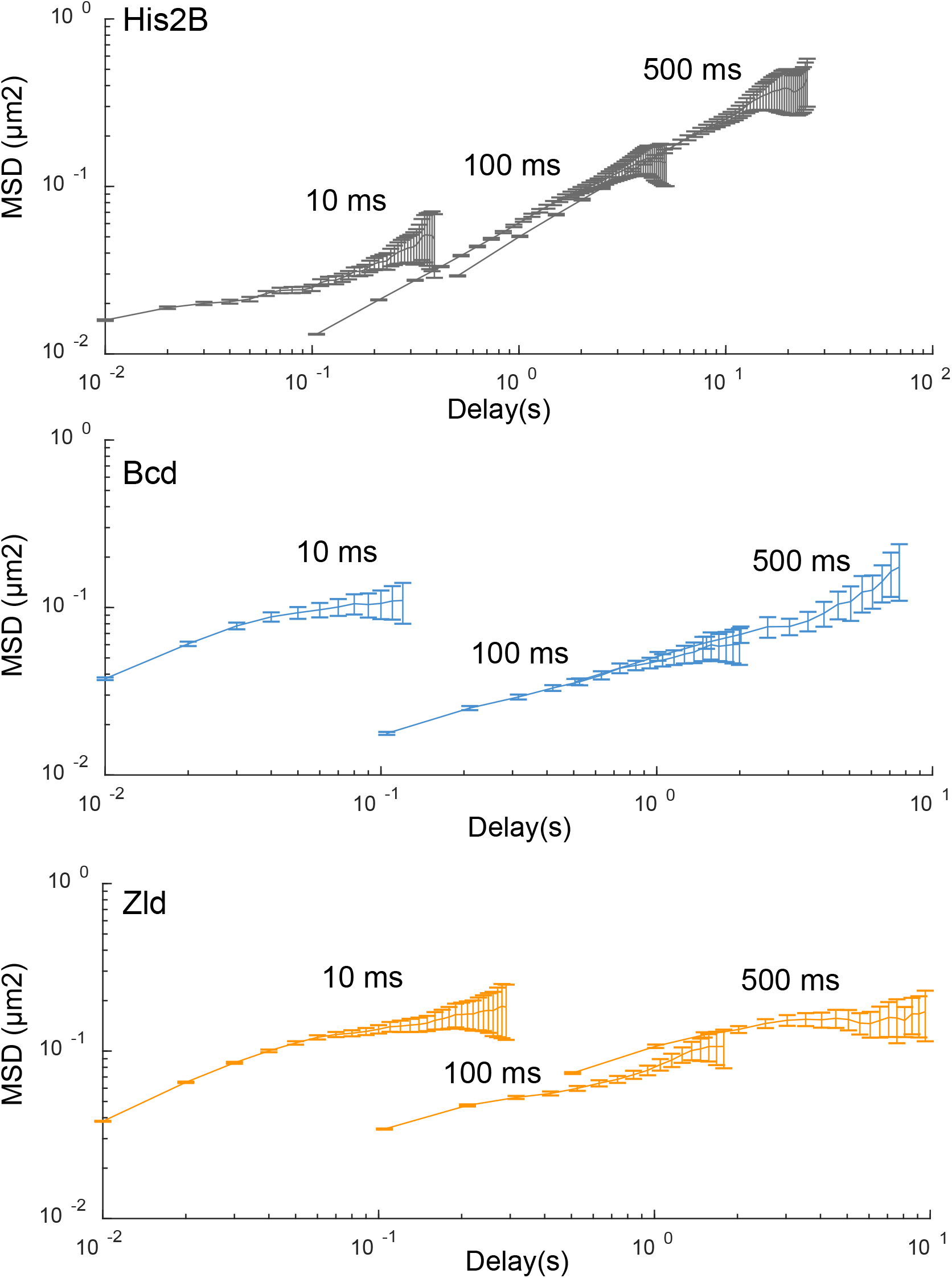
Mean square displacement (MSD) curves. MSD curves for His2B, Zld, and BCD, at all frame rates plotted on log-log-scale. Calculated over 77869, 81660, 11003, at 10 ms, 107998, 42698, 8990 at 100 ms, and 2420, 14487, 47681 at 500 ms, trajectories for His2B, Zld, and Bcd respectively. Error bars show standard error over all trajectories.

To quantify genome average residence times we fit the 500 ms survival probability distributions for Bcd and Zld to a two-exponential decay model (Figure 4B) to estimate the time constants associated with shortand longbinding events. As has been shown previously, the slow and fast time constants associated with the two exponents can be interpreted as the off-rates associated with non-specific and specific binding respectively (Hansen et al. 2017; Mir et al. 2017; Teves et al. 2016; J. Chen et al. 2014). The resulting fits for Zld and Bcd are then bias corrected for photo-bleaching and defocalization using the fits to the His2B data (Hansen et al. 2017).

Using this approach, we estimate genome average residence times for the specifically bound populations of ~5 sec and ~2 sec for Zld and Bcd respectively (Figure 4C). This estimate for Bcd is slightly higher than we obtained previously (Mir et al. 2017), which we attribute to the more accurate bleaching correction using His2B here. These residence time estimates are consistent with FRAP measurements (Figure 2-figure supplement 1) where we measure recovery half times of ~5 and ~1 sec for Zld and Bcd respectively.

Finally, prompted by a visual comparison of the Zld and Bcd trajectories with those of His2B and the relatively high diffusion coefficients for the bound population of all three proteins (Figure 3- figure supplement 1), we explore in more depth the kinetics of molecules that are relatively immobile to the extent that they don’t motion blur into the background at 500 ms exposure times. We thus calculated and compared the time and ensemble averaged mean square displacement (TAMSD) of all three proteins (Figure 4- figure supplement 2). While TAMSD is not an appropriate metric for quantifying diffusion coefficients and bound fractions when the data contain a mixture of different dynamic populations such as at the 10 ms frame rate data (Izeddin et al. 2014; Kepten et al. 2015; Hansen et al. 2018), we reasoned that it is appropriate for a qualitative evaluation of the trajectories from the 500 ms data where we are measuring relatively stable immobile populations.

As expected for transcription factors and proteins confined within an environment, the TAMSDs for all three protein scale as ~⊤^α^, where tau is the lag time and α is the anomalous exponent, consistent with anomalous or sub-diffusive motion, (Normanno et al. 2015; Miné-Hattab et al. 2017; Izeddin et al. 2014). To assess the level of anomalous motion we plotted the TAMSD/⊤ curves from the 500 ms data for all three proteins in loglog scale (Figure 4D). Plotted in this manner a population of molecules exhibiting completely free diffusion would exhibit a log(TAMSD/⊤) curve of slope 0, that is when α=1, whereas sub-diffusive population (0<α<1), display higher negative slopes, with lower α corresponding to more anomalous or confined motion. Strikingly, while Bcd at 500 ms has a high α value similar to His2B, Zld has a low α value consistent with a high amount of anomalous motion (Figure 4D).

Anomalous or subdiffusive motion can result from a range of underlying physical interactions including aggregation, weak interactions with other proteins and chromatin, repetitive binding at proximal binding sites among many other models (Woringer and Darzacq 2018). The complexity of the TAMSDs from Zelda trajectories acquired at 500 ms suggest that at these frame rates we likely measure a mixture of effects that lead to a relatively immobile population of Zelda. Given that Zld is known to exhibit an extremely heterogeneous subnuclear spatial distribution we therefore next examined the bulk rather than single molecule spatialtemporal dynamics of Zld.

### Zelda and Bcd form dynamic subnuclear hubs

Recently, a highly clustered spatial distribution of Zld was reported using confocal microscopy (Dufourt et al. 2018), but the temporal dynamics of these clusters have not been examined due to the technical limitations of confocal microscopy. We thus performed high resolution 4D imaging using LLSM of Zld in developing embryos. We find that the spatial distribution of Zelda is highly dynamic and linked to the nuclear cycle (Video 10). We observe that Zld rapidly loads into nuclei near the end of telophase and associates to the still condensed chromatin. As the chromatin decondenses and the nuclei enter interphase Zld breaks into smaller highly dynamic clusters (Figure 5A and Video 11). As the nucleus enters prophase and the nuclear membrane begins to break up, Zld appears to leave the nucleus and correspondingly the cytoplasmic signal around the nucleus increases (Video 10). As the Zld concentration around the chromatin drops, so does the appearance of clusters, though Zld appears to remain associated with chromatin until the end of prophase. From the end of prophase to telophase no Zld is observed in proximity of the condensed chromatin until it rapidly loads back in to reforming nuclei at the end of telophase.

**Figure 5.**
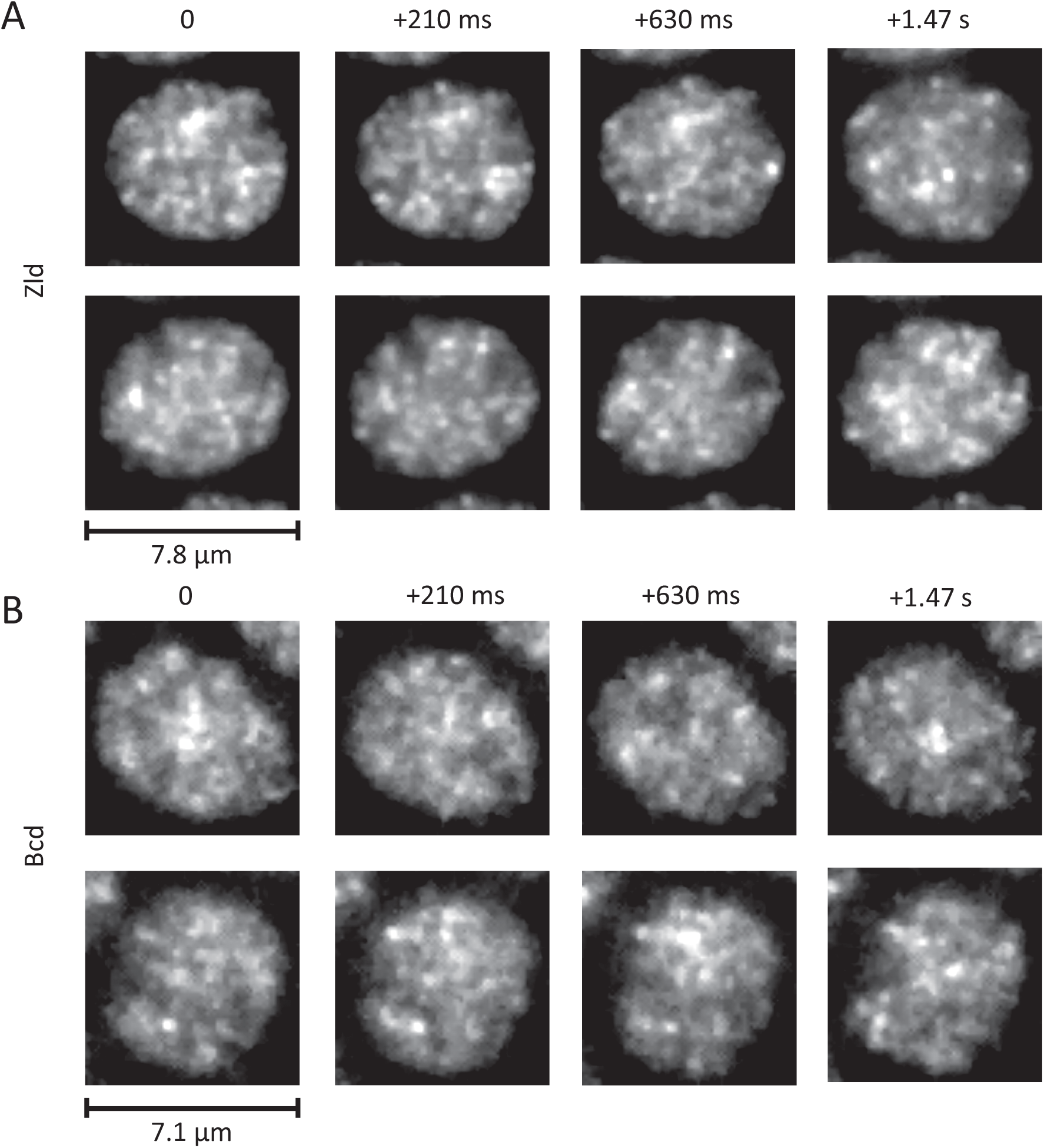
Dynamic interphase hubs of Zld and Bcd. Examples images of the spatial distributions of Zld **(A)** and Bcd **(B)** at various time intervals illustrating the dynamic nature and wide range of size distributions and temporal persistences of enriched hubs (Also see Videos 11-13). mNeonGreen-Zld and EGFP-Bcd were imaged at 15 ms and 210 ms frame rates respectively. To allow comparison, the sum projection of 14 frames (210 ms total integration) is shown for Zld. Images were processed with a 1-pixel radius median filter to remove salt-and-pepper noise and contrast-adjusted manually for visual presentation.

**Figure 3 - figure supplement 1.**
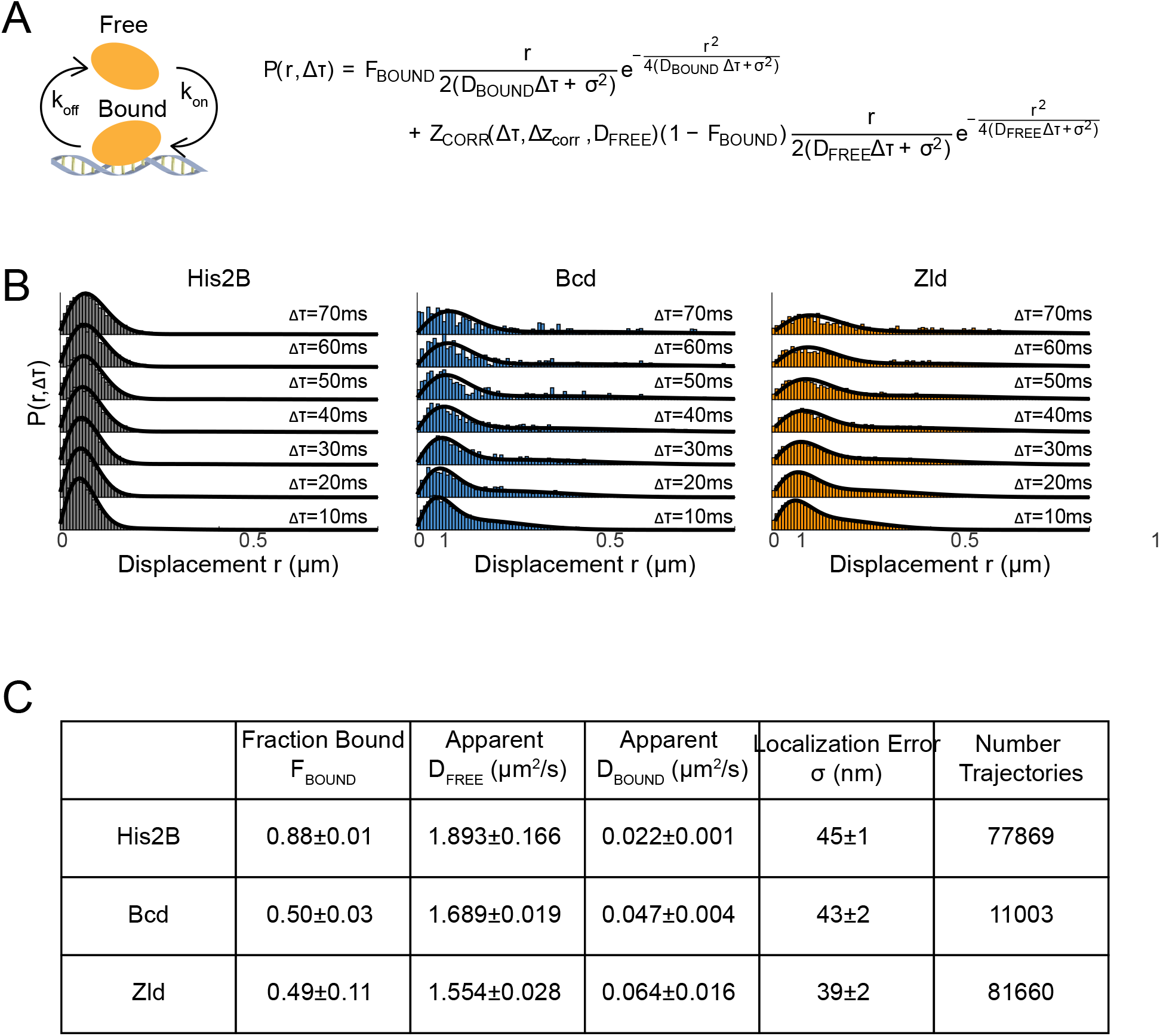
Kinetic modelling of fast SPT data. **(A)** Overview of 2-state model in which molecules are either in a free or bound state and the kinetic-model used to fit the displacement distributions, P(r,Δ⊤), where FBOUND is the fraction of molecules bound, r is the displacement, Δ⊤ is the time delay, σ is the localization error, and ZCORR is the function used to estimate the fraction of trajectories lost due to moving out of the axial detection range (Δzcorr) as detailed in Hansen et al., 2018. **(B)** Displacement distributions at multiple time delays and corresponding fits (black lines), data from 1 embryo is shown for each protein, 3 embryos were measured for each in total. **(C)** Summary of results from fits to the kinetic model where errors shown are over 3 biological replicates.

The dynamic interphase hubs of Zld appear to have a wide distribution of sizes and persist for highly variable amounts of time (Figure 5A and Video 11). When imaged at higher temporal resolutions (Video 12) we observe that both hub location and intensity vary even at subsecond time-scales, suggesting that there is dynamic exchange of Zld molecules in clusters with the rest of the nucleoplasm. Bulk imaging of Bcd also reveals that it forms dynamic hubs in interphase (Figure 5B and Video 13) although they appear less prominent both in size and temporal persistence than those of Zld. Our observations of the nuclear cycle dynamics of Bcd are consistent with previous reports of it filling into the nucleus after mitosis (slower than Zld) and a slow decrease in concentration after the nuclear membrane breaks down (Gregor et al. 2007).

These observations of highly heterogeneous and dynamic sub-nuclear distributions are consistent with our earlier work where we observed that Bcd binding is clustered in discrete subnuclear hubs (Mir et al. 2017). We also previously showed that these Bcd hubs do not form in the absence of maternal Zld which naturally led us to next ask whether there is a relationship between Bcd binding and the local concentration of Zld.

### Bicoid binding events are enriched in Zelda hubs

To explore the relationship between Zld hubs and Bcd binding, we performed dual-color experiments recording the single molecule dynamics of Bcd using mEos3.2Bcd and the bulk spatial distribution of Zld using mNeonGreen-Zld. To strike a balance between the constraints of the imaging system, the dynamic range of the single molecule trajectories, and the fast dynamics of Zld hubs (Figure 5 and Figure 6A) we acquired a bulk fluorescence image of mNeonGreen-Zld with a 1 sec acquisition time followed by 10 frames of single molecule images with a frame rate of 100 ms (Videos 14).

Using the bulk Zld data we partitioned nuclei into regions of high and low Zld density (Figure 6B-C and Figure 6-figure supplement 1). Parsing the Bcd single-molecule data we find that the density of bound Bcd molecules is consistently higher within the Zld enriched regions (Figure 6D). In the embryo anterior, where Bcd concentrations are highest, there is a two-fold increase in the density of Bcd trajectories in high density Zld regions compared to the rest of the nucleoplasm. The density of Bcd trajectories within the enriched Zld regions increases along the anteroposterior axis of the embryo as the Bcd concentration decreases to an excess of around four-fold in the posterior (Figure 6D). This observation is consistent with our previous report of more pronounced clustering of Bcd in the posterior embryo (Mir et al. 2017). When we examined the stability of Bcd binding as a function of Zld enrichment, we also find that at more posterior embryonic positions longer binding events of Bcd are associated with higher Zld enrichment in contrast to the embryo anterior (Figure 6- figure supplement 2). We note that while there is an increase in long Bcd binding events in Zld hubs this effect is less pronounced than the overall enrichment of all Bcd binding events, suggesting that Zld increases the time averaged Bcd occupancy at DNA binding sites by increasing its local concentration (increasing *k*_*on*_) and not by altering its residence times at its target sites (increasing *k*_*off*_).

**Figure 6.**
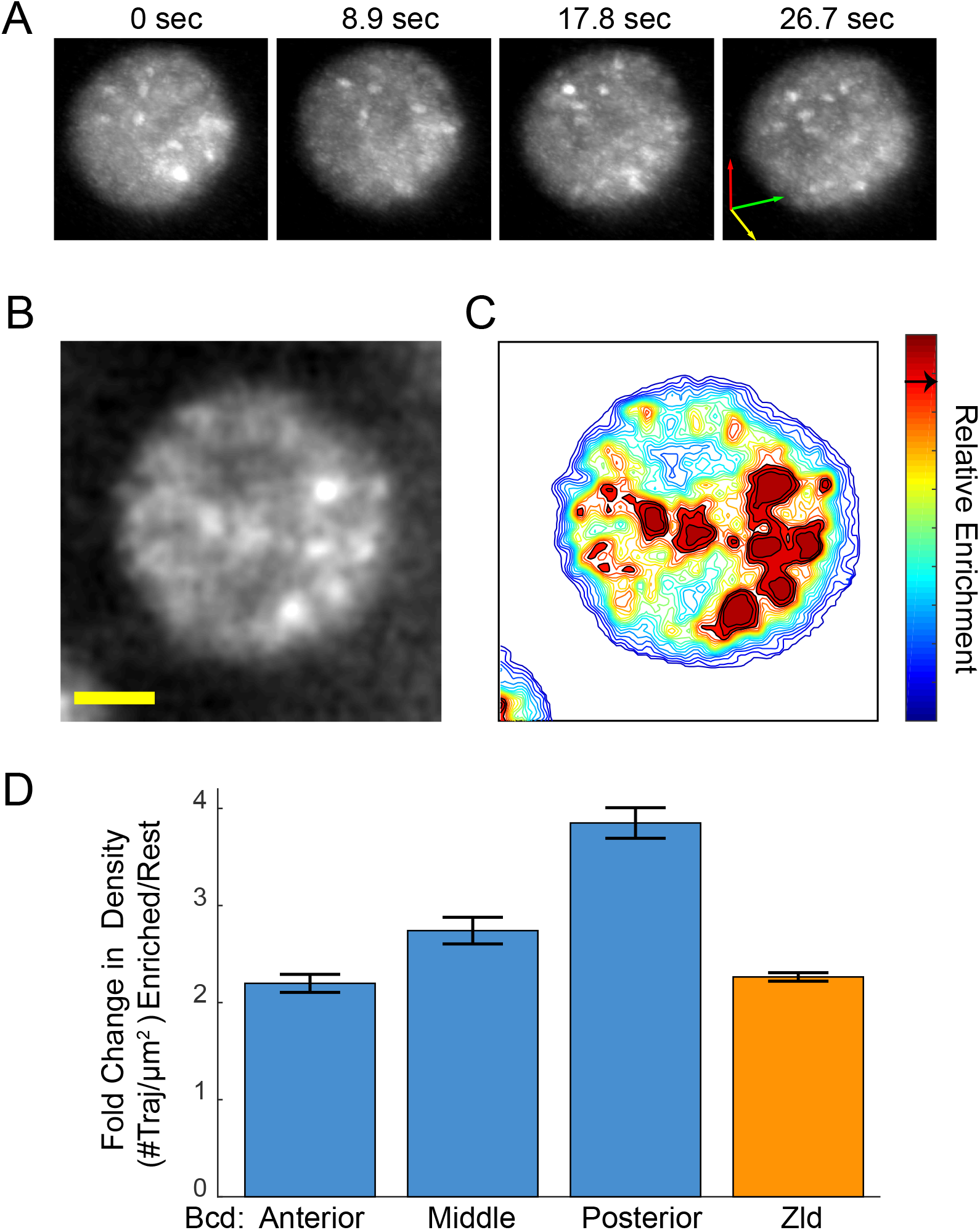
Enrichment and immobilization of Bcd within Zld hubs. **(A)** Three-dimensional volume renderings of an interphase nucleus showing the dynamic nature of Zld hubs (see Videos 11-12). The 3D axes indicate the xyz axes and the arrow lengths are 2 μm along each direction. **(B)** Representative snapshot of the interphase distribution of Zelda, yellow scale bar is 2 μm. **(C)** Relative enrichment map for the nucleus shown in (B), the arrow on the colorbar indicates the threshold for defining a region as enriched. **(D)** Fold change in density of single molecule trajectories of Bcd (across anteroposterior axis) and Zld in Zld enriched regions vs. the rest of the nucleoplasm. Error bars show standard error over 3 embryos with a total of 1344, 3921, 481 nuclear images for Bcd Ant, Mid, and Post respectively, and 4399 for Zld.

**Figure 6 - figure supplement 1.**
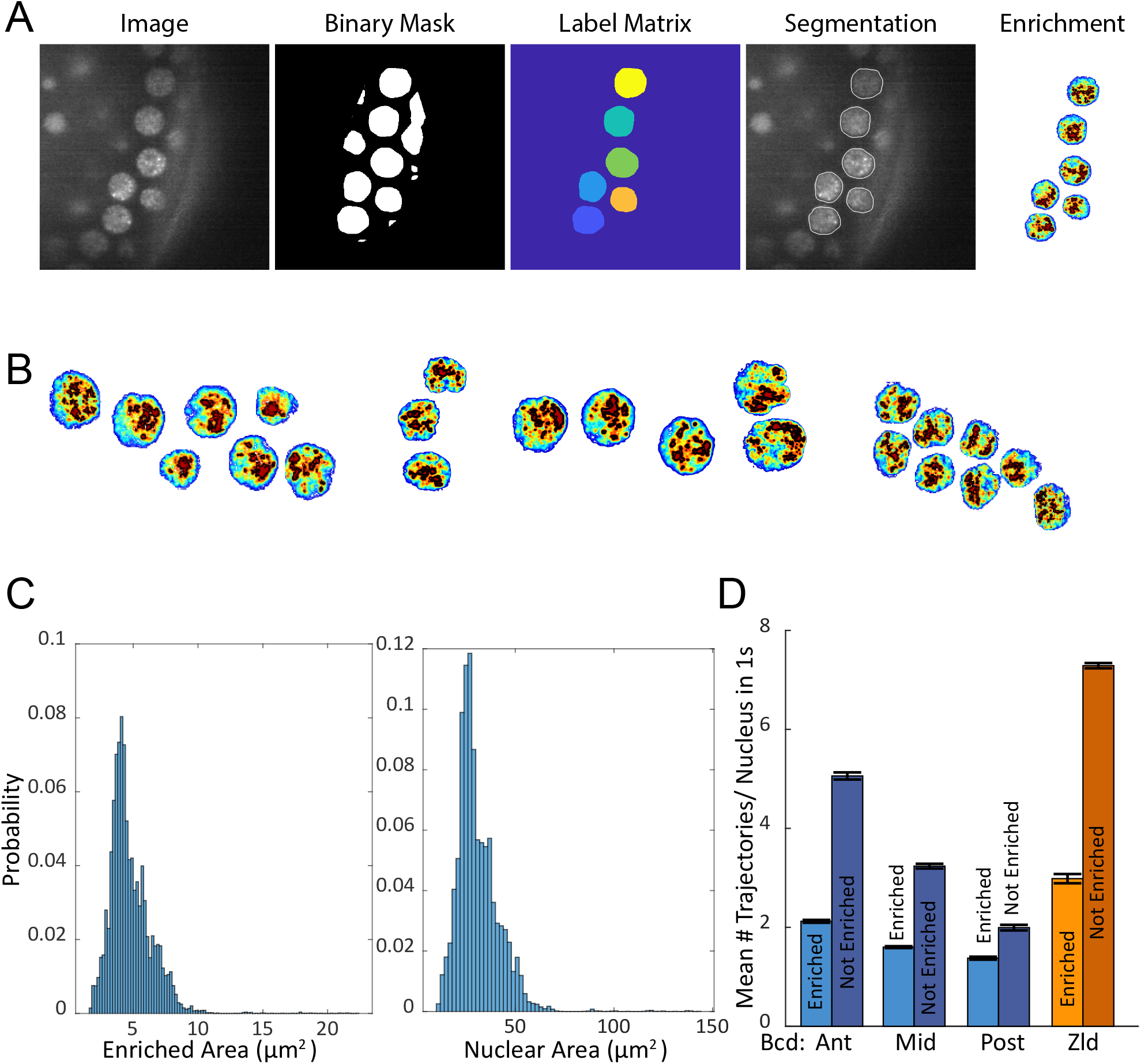
Analysis of Zelda enrichment. **(A)** Major steps in nuclear segmentation algorithm, the Zelda image is first low pass filtered and then converted to a binary mask through adaptive thresholding, a hand drawn mask is applied to remove regions of low contrast and the embryo edge. The binary mask is dilated, a label matrix is calculated and regions not meeting eccentricity and size cutoffs are removed. The relative enrichment map for each nucleus is then calculated using the distribution of intensity values within each nucleoplasm. **(B)** More examples of calculated enrichment maps. **(C)** Histograms of enriched areas and nuclear areas. Distributions are from 33041 images of nuclei. **(D)** Mean number of single trajectories in enriched vs. not enriched regions per nuclear image, for a total of 1344, 3921, 489, nuclear images in the Anterior (Ant), Middle (Mid), and Posterior(Mid) positions for Bcd respectively, and 4399 for Zld with trajectories (#Enriched/#Not) of 4326/12437, 3921/11211, 836/1793 for Bcd Ant, Mid, and Post respectively and 25470/ 66923 for Zld. Error bars are standard errors over all nuclear images.

**Figure 6 - figure supplement 2.**
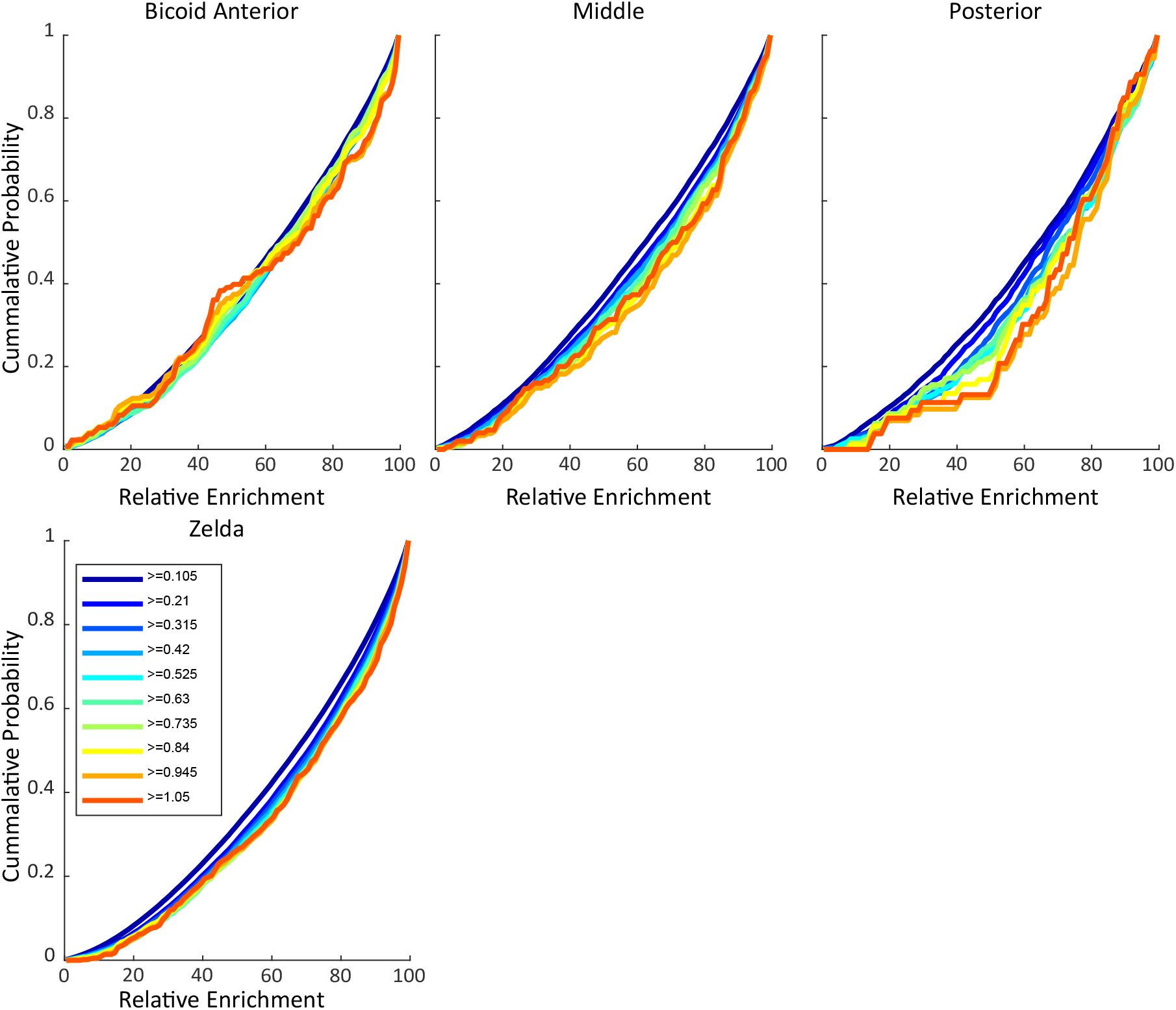
Cumulative probability of trajectories vs. relative enrichment. Plots show the cumulative probability of a trajectory greater than equal to a certain length (as indicated by color) vs. the relative enrichment of Zld. The top row shows the probability for single molecule trajectories of Bicoid across the Anteroposterior axis and the bottom panel is for Zld single molecule trajectories. For Bcd binding events, In the anterior there is little cost in terms of enrichment of increased trajectory lengths but in middle and posterior positions greater Zld enrichment is required for molecules to be immobilized for longer.

This association between Zld hubs and Bcd binding suggests that these hubs, though dynamic and transient, might be preferentially forming on genes that are coregulated by Zld and Bcd. Furthermore, given the strong association of Zld binding measured by chromatin immunoprecipitation with the binding of many early embryonic factors (Harrison et al. 2011) we expected a strong correlation between Zld hubs and sites of active Bcd-dependent transcription. To test this hypothesis we next performed imaging of the spatial distributions of Bcd and Zld in the context of active transcription.

### Zelda hubs are not stably associated with sites of active transcription

We chose to study the relationship between Zld and Bcd hubs and transcriptional activity at the canonical Bcd target gene *hunchback* (*hb*). The *hb* gene was the first identified target of Bcd (Struhl, Struhl, and Macdonald 1989; Tautz 1988), and its anterior transcription is dramatically disrupted in the absence of Bcd (Staller et al. 2015; Ochoa-Espinosa et al. 2009; Hannon, Blythe, and Wieschaus 2017). The regulatory sequences for *hb* contain multiple clustered Bcd binding sites, as well as recognizable Zelda motifs (Harrison et al. 2011) (Figure 7-figure supplement 1). ChIP studies show that both Zelda and Bcd bind strongly at the *hb* locus, though loss of Zelda has only a modest quantitative effect on *hb* expression (Combs and Eisen 2017; Nien et al. 2011). An enrichment of Bcd in the vicinity of active *hb* loci was previously observed using FISH (H. Xu et al. 2015) on fixed embryos, but nothing is known about the dynamics of this enrichment or its relationship to Zld.

**Figure 7.**
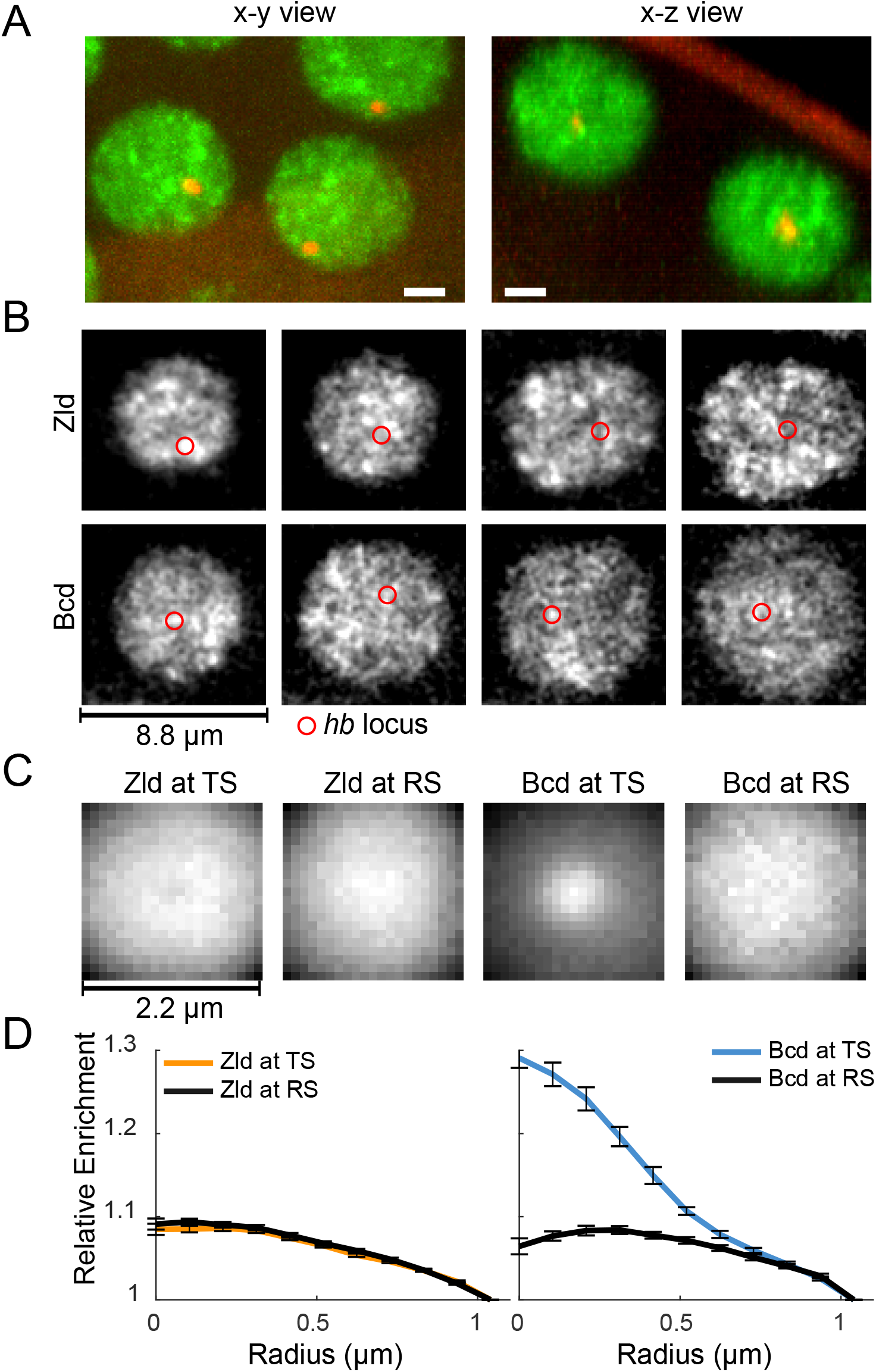
Spatio-temporal distribution of Zld and Bcd hubs in context of active *hb* loci. **(A)** Representative x-y and x-z max projections over a nuclear diameter of mNeonGreen-Zld (green) and an active hb locus tagged with MS2-MCP-mCherry (red) white scale bars are 2 μm **(B)** Representative snapshots of the distribution of Zld and Bcd with the *hb* locus indicated by the red circle. Images suggest that high concentration Bcd hubs frequent the active locus whereas Zld exhibits more transient and peripheral interactions. Contrast of each image was manually adjusted for visualization and comparison. **(C)** Average Zld and Bcd signals in a 2.2 μm window centered at active hb locus (TS) and at random sites in the nucleus (RS). Averages were calculated over 3943 and 6307 images of active loci from 6 embryos for Bcd and Zld respectively (see Videos 18 and 19). **(D)** Radial profiles of the images in C, normalized to 1 at the largest radius. Error bars show standard error over all images analyzed.

**Figure 7 - figure supplement 1.**
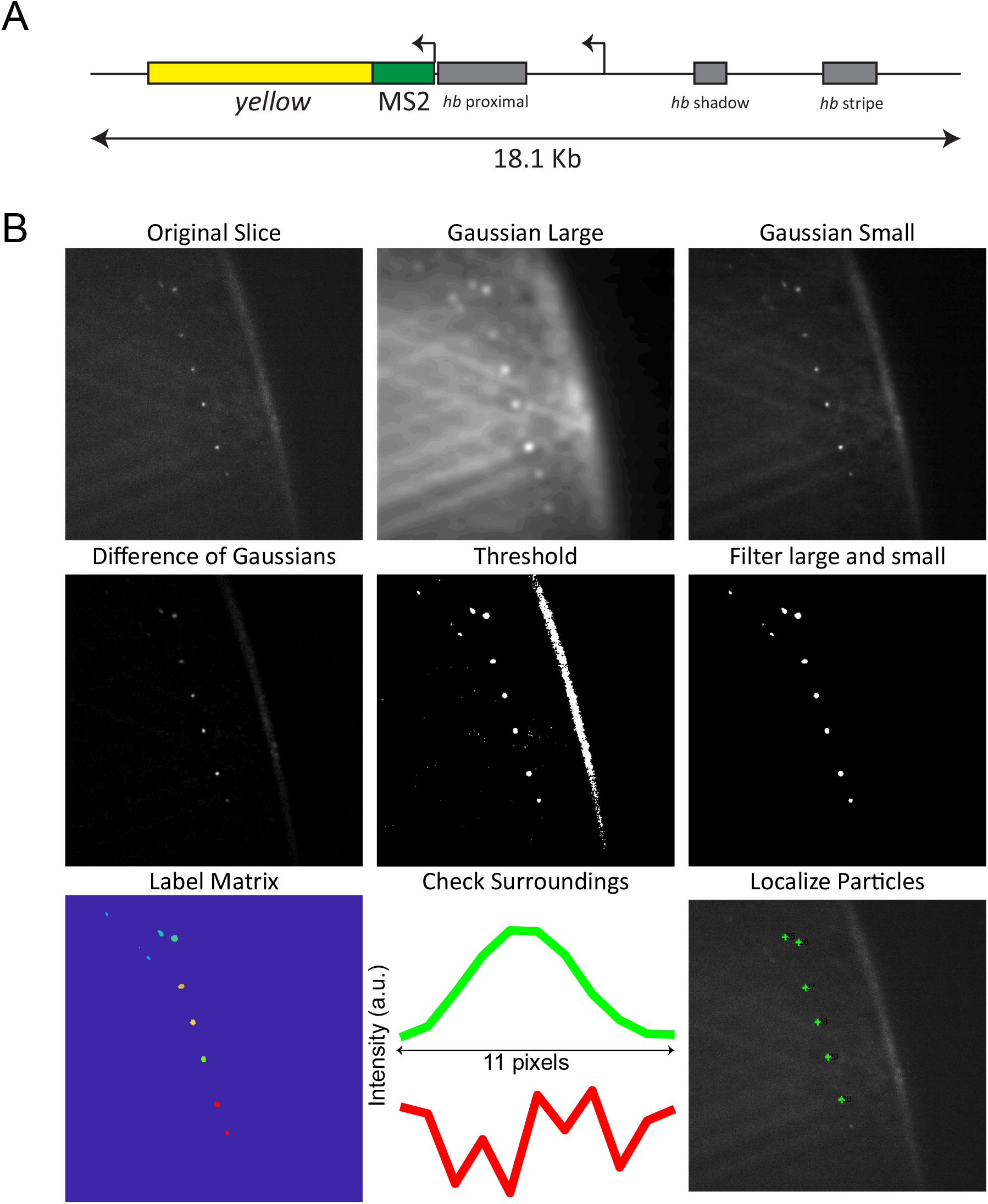
MS2 system and data analysis. **(A)** BAC construct used in the experiments to visualize transcription of hb, from (Bothman et al. 2015). The coding region is replaced by the yellow gene and 24 MS2 stem loops are inserted downstream of the P2 promoter, allowing the visualization of transcripts from both proximal and distal promoters. The BAC spans an 18.1 kb genomic region containing all known hb regulatory sequences, and its expression closely mirrors that of native hb. **(B)** Segmentation and localization of active loci. A difference of Gaussians is used to enhance the contrast of transcription sites, a binary mask is calculated by applying a threshold and filtered to remove large and small structures. A label matrix for remaining segmented regions is then generated and line profiles through the center of the regions is used to determine if they lie inside of a nucleus. Finally, the center of the active locus is determined using the line profile.

To visualize the *hb* locus, we took advantage of the MS2 system, which allows fluorescent labelling of nascent transcripts of specific genes (Garcia et al. 2013; Bothma et al. 2015). Bothma, et al. generated a fly line carrying a bacterial artificial chromosome (BAC) that contains an MS2-labeled *hb* locus that closely recapitulates the expression of *hb* itself (Bothma et al. 2015). We thus performed high spatio-temporal resolution 4D imaging of the bulk distributions of Zld, and separately Bcd, in embryos carrying the *hb* BAC and MCP-mCherry (Figure 7A, Figure 7-figure supplement 1 and Videos 15-17).

From visual examination of movies of Zld and Bcd in the presence of *hb* transcription (Videos 16-17 and Figure 7B), we observe that the temporal relationship between Zld and Bcd hubs and the *hb* locus in nuclei where it is expressed (and therefore visible) is highly dynamic. We do not observe stable associations between high-concentration hubs of either factor and *hb*. However, we do see that contacts between *hb* and hubs of both factors occur frequently, so we next asked whether *hb* showed any preferential association with hubs of either factor over time.

Following (Spiluttini et al. 2010), we averaged the Bcd and Zld signal surrounding active *hb* loci over thousands of images from 6 embryos (Figure 7C and Videos 18-19). For Bcd, we observe a sharp enrichment of fluorescent signal at the *hb* locus in comparison to randomly selected control points within nuclei (Figure 7D). We observe no such enrichment for Zld at *hb* (Figure 7B), however we note that Zld has many fold more targets than Bcd (Harrison et al. 2011; Li et al. 2008), and its target loci may be at too high a density in the nucleoplasm to detect enrichment at any one of them with this assay. These results imply a previously unappreciated aspect of the relationship between transcription factor hubs and their target genes: that individual hubs are multifactorial and likely service many different genes and loci within the nucleus.

## Discussion

We previously reported a strong correlation between genomic locations of Zld and Bcd binding (Harrison et al. 2011) and suggested, based on a roughly twenty fold increase in the occupancy of potential Bcd binding sites in Zld bound regions, that there is strong cooperativity between these two factors. More recently we showed that Bcd binds DNA highly transiently, but that its binding is spatially organized in a Zld dependent manner (Mir et al. 2017).

Based on these data we hypothesized that Zld could act as a DNA bound scaffold facilitating Bcd binding by increasing its local concentration in the vicinity of its target. Here, however, we find that Zld also binds DNA transiently and therefore cannot, by itself, act as a stable scaffold at enhancers.

Our observation that Zld and Bcd form hubs of locally high concentration suggests an alternative model in which multiple factors enriched within hubs interact to increase factor occupancy at DNA targets without the need for stabilizing this interaction by conventional “lock and key” interactions. In our model efficient occupancy of a site is achieved by frequent transient weak binding events within hubs rather than long stable interactions on DNA (Woringer and Darzacq 2018).

A strong correlation between transcription and hubs of RNA polymerase II (Cisse et al. 2013; Chong et al. 2018; Boehning et al. 2018) and transcription factors (J. Chen et al. 2014; Chong et al. 2018; Wollman et al. 2017; Liu et al. 2014) has been reported in mammalian cells. In *Drosophila*, high local concentrations of the transcription factor Ultrabithorax (Ubx) at sites of Ubx mediated transcription has recently been reported (Tsai et al. 2017).

However, use of LLSM to image hubs at high frame rates shows that they are not stable structures, and furthermore that they interact only transiently with sites of active transcription. We suggest that what previous reports have described are actually time-averaged accumulations (such as what we observe for Bcd at *hb*) rather than discrete sub-nuclear bodies.

The most surprising observation we report is that at an active locus, transcription occurs with rare and transient visits of hubs containing the primary activator of the locus. One possible explanation is some version of the decades old “hit-and-run” model proposed by (Schaffner 1988) in which transcription factor binding and interactions with enhancers are only required to switch promoters into an active state, after which multiple rounds of transcription could occur in the absence of transcription factor binding (Para et al. 2014; Doidy et al. 2016). Transcription could then be regulated by the frequency of visits, rather than stable association.

Our observation that hubs form rapidly at the exit from mitosis and are most pronounced at times when no transcription happens raises the possibility of an even greater temporal disconnect. We and others have suggested that a primary role of Zld is in the licensing of enhancer and promoter chromatin for the binding of other factors (Li et al. 2014; Foo et al. 2014; Schulz et al. 2015; Sun et al. 2015) and in the creation and stabilization of chromatin three-dimensional structure (Hug et al. 2017). It is possible that the key point of activity occurs early during each nuclear cycle when the chromatin topology is challenged by the replication machinery and transcription has not begun (Blythe and Wieschaus 2016).

Our single-molecule data point to intrinsic forces that might lead to the formation and maintenance of Zld hubs. The highly anomalous nature of the movement of Zld that differs from that of chromatin associated His2B and Bcd suggests that its motion depends at least in part on interactions off of DNA. Most of the amino acid sequence of Zld consists of intrinsically disordered domains, some of which are required for its function (Hamm et al. 2017). We and others have shown that intrinsically disordered domains mediate weak, multivalent protein-protein interactions between regulatory factors (Chong et al. 2018; Boehning et al. 2018; Kovar 2011; Burke et al. 2015; Altmeyer et al. 2015; Xiang et al. 2015; Friedman et al. 2007) that lead to the selective enrichment of factors in hubs (Chong et al. 2018; Boehning et al. 2018). We think it is therefore highly likely that Zld, and therefore Bcd, hubs are formed by interactions involving intrinsically disordered domains.

Bcd is only one of many proteins whose early embryonic binding sites have a high degree of overlap with those of Zld (Harrison et al. 2011), including many other factors involved in anterior-posterior and dorsal-ventral patterning, and other processes (Reichardt et al. 2018; Foo et al. 2014; Sun et al. 2015; Schulz et al. 2015; Nien et al. 2011; Z. Xu et al. 2014; Pearson, Watson, and Crews 2012; Boija and Mannervik 2016; Shin and Hong 2016; Ozdemir et al. 2014). We hypothesize that Zld provides scaffolds to form distinct hubs with each of these factors, mediated by a combination of weak and transient protein:protein and protein:DNA interactions.

Such hubs could help explain the disconnect between the canonical view of transcription factor function based on their directly mediating interactions between DNA and the core transcriptional machinery and data like those presented here that suggests a more stochastic temporal relationship. There is abundant evidence that transcription factors can affect promoter activity by recruiting additional transcription factors, chromatin remodelers and modifiers, and other proteins. Extrapolating from our observation that Zld appears to form some type of scaffold for Bcd hubs, we propose that hubs contain not only transcription factors, but loose assemblages of multiple proteins with diverse activities. Such multifactor hubs could provide each transcription factor with a bespoke proteome, with far greater regulatory capacity and precision than could plausibly be achieved through stable direct protein-protein interactions involving each factor.

## Video Legends

**Video 1 - Movie illustrating ability to controllably photactiviate mEOS3.2**

Related to Figure 1. Movie illustrating the ability to photo-activate mEos3.2 with the modified LLSM system. The power of the 405 nm laser was set to its highest value at approximately the 4 second mark after which a continuous increase in the signal can be observed reflecting the density of activated molecules.

**Video 2 - Example movie of mEos3.2-Zld acquired at 10 ms frame rate**

Related to Figure 1. White scale bar is 5 μm, images were gaussian filtered and inverted for display.

**Video 3 - Example movie of mEos3.2-Zld (red) and His2B-EGFP (green) acquired at 100 ms frame rate**

Related to Figure 1. White scale bar is 5 μm, a nucleus undergoing division is shown for illustrative purposes, data from mitotic nuclei were not used for single molecule analysis in this work. Images were gaussian filtered for display.

**Video 4 - Example movie of mEos3.2-Zld (red) and His2B-EGFP (green) acquired at 500 ms frame rate**

Related to Figure 1. White scale bar is 5 μm, data from mitotic nuclei were not used for single molecule analysis in this work. Images were gaussian filtered for display.

**Video 5 - Example of a mobile molecule of mEos3.2-Zld tracked at 10 ms frame rate**

Related to Figure 1. White scale bar is 2 μm. Images were gaussian filtered and inverted for display.

**Video 6 - Example of a immobile molecule of mEos3.2-Zld tracked at 10ms frame rate**

Related to Figure 1. White scale bar is 2 μm. Images were gaussian filtered and inverted for display.

**Video 7 - Comparison of single molecule movies for His2B-mEos3.2, mEos3.2-Bcd, and mEos3.2-Zld at 10 ms frame rate**

Related to Figure 2. White scale bar is 2 μm. Panels left to right are His2B-mEos3.2, mEos3.2-Bcd, and mEos3.2-Zld. Images were gaussian filtered, inverted, and contrast was manually adjusted to allow for ease of visual comparison.

**Video 8 - Comparison of single molecule movies for His2B-mEos3.2, mEos3.2-Bcd, and mEos3.2-Zld at 100 ms frame rate**

Related to Figure 2. White scale bar is 2 μm. Panels left to right are His2B-mEos3.2, mEos3.2-Bcd, and mEos3.2-Zld. Images were gaussian filtered, inverted, and contrast was manually adjusted to allow for ease of visual comparison.

**Video 9 - Comparison of single molecule movies for His2B-mEos3.2, mEos3.2-Bcd, and mEos3.2-Zld at 500 ms frame rate**

Related to Figure 2. White scale bar is 2 μm. Panels left to right are His2B-mEos3.2, mEos3.2-Bcd, and mEos3.2-Zld. Images were gaussian filtered, inverted, and contrast was manually adjusted to allow for ease of visual comparison.

**Video 10 - Cell Cycle Dynamics of Zelda spatial distribution**.

Related to Figure 5. White scale bar is 2 μm. Panels left to right are His2B-mEos3.2, mEos3.2-Bcd, and mEos3.2-Zld. Maximum intensity projection over 81 slices spaced at 250 nm apart. Right His2B-mCherry left sfGFP-Zld showing through nuclear cycles 13 and 14. His2B is present at high concentrations in both the cytoplasm and nucleoplasm of Drosophila embryos, resulting in the extranuclear signal visible in the right panel. Time interval between each volume is 8.91 s. The field of view is 23.2×23.2 μm.

**Video 11 - Four dimensional Interphase dynamics of Zld spatial distributions**

Related to Figure 5. Three-dimensional rendering of Zld spatial distribution in nuclear cycle 13. Progress bar indicates time in minutes. Slice spacing in z is 200 nm, time between volumes is 9.5 s.

**Video 12 - Interphase dynamics of Zld spatial distributions at high temporal resolution**

Related to Figure 5. Interphase dynamics of Zld spatial distribution at 10 ms exposure times and 15 ms frame rate.

**Video 13 - Four dimensional dynamics of Bcd spatial distribution**

Related to Figure 5. Three-dimensional rendering of Bcd spatial distribution from nuclear cycle 13-14. Progress bar indicates time in minutes. Slice spacing in z is 200 nm, time between volumes is 8 s.

**Video 14 - Bcd single molecule localizations in context of the bulk spatial distribution of Zld**

Related to Figure 6. Bicoid detections (red dots) overlaid on Zld spatial distribution. Each frame in the movie corresponds to 1 s of detections at 100 ms exposure times. Field of view is 9 x 14 μm.

**Video 15 - Four dimensional imaging of protein distribution in the context of transcription**.

Related to Figure 7. Example of four dimensional imaging of Zld spatial distribution in the context of an active transcription site imaged using the MS2 system (red).

**Video 16 - Example of Bcd spatial distribution around an active *hb* locus**.

Related to Figure 7. eGFP-Bcd (green) and MCP-mcherry (red) dynamics from nuclear cycles 13-14. Maximum intensity projection over 61 slices spaced 250 nm apart with ~6 s between volumes.

**Video 17 - Example of Zld spatial distribution around an active *hb* locus**.

Related to Figure 7. mNeonGreen-Zld (green) and MCP-mcherry (red) dynamics from nuclear cycles 13-14. Maximum intensity projection over 61 slices spaced 250 nm apart with 5.18 s between frames.

**Video 18 - Calculation of average Zld signal around active *hb* loci**

Related to Figure 7. Top panel shows images of the MS2 signal, corresponding Zld signal in the same window (TS) and at a random control site in the same nucleus (RS). Middle panel shows a running average of the images in the top panel and bottom panel shows the corresponding running average radial profile.

**Video 19 - Calculation of average Bcd signal around active *hb* loci**

Related to Figure 7. Top panel shows images of the MS2 signal, corresponding Bcd signal in the same window (TS) and at a random control site in the same nucleus (RS). Middle panel shows a running average of the images in the top panel and bottom panel shows the corresponding running average radial profile.

## Methods

### Generation of transgenic fly lines

The following fly lines were constructed using CRISPR/Cas9 mutagenesis with homology directed repair: mNeonGreen-Zelda, mEos3.2-Zelda, mEos3.2-Bicoid. sgRNAs targeting sites near the desired insertion sites were cloned via the primer annealing method into plasmid pMRS-1, which is a version of pCFD3 (Port et al. 2014) (addgene #49410) with alterations to the sgRNA body made according to (B. Chen et al. 2013). Homology directed repair templates were constructed in a pUC19 backbone via Gibson assembly with the desired tag, with an N-terminal FLAG tag, flanked by 1 kb homology arms. We tested a number of linker sequences and found variable tag- and protein-specific effects on viability. Linker sequences that yielded homozygous viable animals were GDGAGLIN (mNeonGreen-Zld), GGGGSGSGGS (mEos3.2-Zld and mEos3.2-Bcd) and GGGGSGSGGSMTRDYKDDDDKTRGS (H2B-mEos3.2 and H2B-EGFP).

HDR template and sgRNA plasmids were sent to Rainbow Transgenic Flies, Inc. (Camarillo, CA) to be injected into embryos expressing Cas9 in the germline. Resulting adult flies were crossed to flies possessing balancer chromosomes matching the relevant chromosome. Single F1 progeny carrying the marked balancer were crossed to balancer stock flies, allowed 4-8 days for females to lay sufficient eggs, and the F1 parents were sacrificed for PCR genotyping. For positive hits, balanced lines were generated by selecting appropriately-marked F2 progeny, and F3 animals were examined for the presence of homozygous animals, revealed by the lack of balancer phenotype. As Bicoid has only maternal phenotypes, homozygous mothers were tested for the ability to give viable offspring. Lines that tolerated homozygous insertions were subjected to further screening by the preparation of clean genomic DNA, amplification of the locus using primers outside the donor homology arms, and subsequent Sanger sequencing of the entire amplicon. Lines carrying insertions free of mutations and containing no incorporated plasmid backbone were kept and utilized for imaging experiments.

HisBmEos3.2 was introduced as a supplemental transgene via PhiC31mediated recombinase (Groth et al. 2004) into landing site VK33 (Venken et al. 2009). A transgene was used to avoid potential complications associated with editing the highly multicopy histone locus.

We chose a red fluorescent protein for single molecule imaging in embryos as better signals are achievable at longer wavelengths. First, as is well known, there is high autofluorescence at greener wavelengths in the *Drosophila* embryo (and for most biological materials). Second, Rayleigh scattering, scattering from particles of sizes less than the wavelength of the imaging light (the phenomenon responsible for blue skies and red sunsets), scales as ~1/λ^4^ where λ is the imaging wavelength. Thus using longer wavelengths results in fewer photons being scattered and thus more photons being absorbed, emitted and collected from single molecules (Mir et al. 2018).

### MS2 crosses

For MS2 experiments, yw; +; MCP-mCherry (gift from S. Alamos and H.G. Garcia) virgin females were crossed to males homozygous for either EGFP-Bcd or mNeonGreen-Zld. Resulting female progeny maternally deposit both MCP and the labeled TF in embryos. Virgin females were crossed to males homozygous for the *hb* MS2 BAC and resulting embryos were used for imaging.

### Lattice LightSheet Microscopy of Live Embryos

For embryo collection a 90 minute laying period in small fly cages. Prior to embryo collection the surface of a 5 mm diameter glass coverslip was made adhesive by deposition of of a small drop of glue solution (the glue solution was prepared by dissolving a roll of double-sided scotch tape in heptane overnight). The coverslip was allowed to dry for at least 5 minutes, which is sufficient time for the heptane to evaporate leaving behind a sticky surface. Embryos were washed off from the cage lids using tap water and gentle agitation with a paintbrush into a nylon cell-strainer basket. Embryos were then dechorionated in 100% bleach for 90 seconds. The dechorionation was then stopped by continuous washing under tap water until no further bleach smell could be detected, typically 30 seconds. The embryos were then transferred from the water filled strainer basket onto an agar pad using a fine haired paintbrush and arranged into an array of typically 3 rows and 5 columns with a consistent anteroposterior (A-P) orientation. The arranged embryos were then gently contact transferred onto the adhesive coverslip which was subsequently loaded into the microscope sample holder.

A home built lattice light-sheet microscope (LLSM) was used (B.-C. Chen et al. 2014; Mir et al. 2017) for all single molecule, bulk fluorescence, and MS2 imaging experiments. Images were acquired using two Hamamatsu ORCA-Flash 4.0 digital CMOS cameras (C13440-20CU). An image splitting longpass dichroic (Semrock FF-560) was placed in between the two cameras to separate emission wavelengths of over and under 560 nm, in addition bandpass filters corresponding to the fluorophore of interest were installed in front of each camera to provide further spectral filtering (Semrock FF01-525/50 for mNeon and sfGFP, Semrock FF01-593/46 for mEOS3.2, and Semrock FF01-629/53 for mCherry). Further details of imaging settings and conditions for each type of imaging experiment are provided in the corresponding sections below.

For all experiments the stage positions corresponding to the anterior and posterior extents of each embryo imaged were recorded. The position along the anteroposterior axis for each image or movie recorded was then calculated as a fraction of the embryonic length (EL) with 0 and 1 to the anterior and posterior extents of the embryo respectively. The nuclear cycle and progression within the nuclear cycle (e.g. interphase, prophase, mitosis) were also recorded for each movie or image. Times between nuclear cycles were also monitored to ensure that data was being acquired on a healthy and normally developing embryos. Embryos which exhibited aberrant development, for example longer than usual nuclear cycles, or numerous aberrant nuclear divisions were abandoned and the data was discarded.

### Single molecule imaging and tracking in live embryos

For single molecule imaging experiments the illumination module of the LLSM was modified to provide constant photo activation using a 405 nm laser line that bypasses the Acousto-optical tunable filter (AOTF) (Figure 1- figure supplement 1). We found that even when a lattice pattern for 561 nm was displayed on the spatial light modulator (SLM) sufficient 405 nm illumination was present in the imaging plane to allow for controlled photo-activation of mEos3.2-Bcd and mEos3.2-Zld. For all single molecule experiments a 30 beam square lattice with 0.55 and 0.44 inner and outer Numerical Apertures respectively was used in dithered mode for excitation. The 405 nm laser line was kept on constantly during the acquisition period for photoswitching and a 561 nm laser line was used for excitation. For both mEos3.2-Bcd and mEos3.2-Zld data was acquired at 7.5, 100 and 500 ms exposure times with effective frame rates of 100, 9.52, and 1.98 Hz respectively. The excitation laser power was optimized empirically for each exposure time to achieve sufficient contrast for single molecule tracking and the powers of the photoswitching laser were also optimized empirically to achieve low enough densities of detections to enable tracking. The excitation laser power was 0.1 mW, 0.6 mW, 2.3 mW and switching laser power was 2.3 µW, 3.9 µW, and 8.5 µW for 500, 100, and 7.5 ms exposures respectively as measured at the back focal plane of the excitation objective. The same settings were used to acquire control data at each exposure time on His2B-mEos3.2. For all exposure times the length of each acquisition was 105 s, corresponding to 200, 1000, and 10000 frames at 500, 100, and 7.5 ms exposure times respectively. The acquisition length was set so that sufficient fields of views could be captured in the short interphase times of the early nuclear cycles while also capturing a sufficient number of single molecule trajectories.

For characterization of single molecule dynamics at these multiple time scales both mEos3.2-Bcd and mEos3.2-Zld were measured in a His2B-EGFP background. The His2B-EGFP channel was used to ensure optimal positioning of the sample within the light sheet, to keep track of progression through a cell cycle, and monitor the development of the embryo. A fortunate bonus was that at 100 ms and 500 ms exposures, there was sufficient excitation of His2B-EGFP from the photoactivation 405 nm laser that we could perform simultaneous imaging of chromatin and single molecule dynamics (Videos 3-4).

For quantification of single molecule mEos3.2-Bcd dynamics in the context of mNeonGreen-Zld, single molecule data was acquired for 1 s (10 frames at 100 ms exposure times), followed by 10 frames of acquisition in the mNeon channel at 10 ms exposure times, and this sequence was then repeated 100 times. The sum of the 10 mNeonGreen images was then calculated to effectively provide a 100 ms exposure image. This scheme was designed such that the dynamic motion of Zld could be captured in addition to the binding kinetics of Bcd with sufficient temporal resolution without having to modify the LLSM control software. The rest of the imaging parameters were kept identical to those described above. For all single molecule experiments nuclei from at least 3 embryos were measured spanning a range of anteroposterior positions and at nuclear cycles ranging from 12-14.

Localization and tracking of single molecules was performed using a MATLAB implementation of the dynamic multiple-target tracing algorithm (Sergé et al. 2008) as previously described (Mir et al. 2017; Hansen et al. 2018, 2017; Teves et al. 2016).

### Mean Square Displacement Analysis

Mean Square Displacement curves were calculated using the open source msdanalyzer package (Tarantino et al. 2014). For analysis of sub-diffusive motion MSD/⊤ curves for His2B, Zld, and BCD plotted on log-log-scale. As for anomalous diffusion MSD(⊤)=Γ⊤^α^, where α is the confinement factor the log(MSD/⊤)=log(Γ) + (α-1)⊤ (Izeddin et al. 2014). The log of the MSD/⊤ was thus used to estimate the range of α values for each protein.

### Analysis of short exposure (10 millisecond) single molecule trajectories

Single molecule trajectories were analyzed using Spot-on (Hansen et al. 2018), a freely available open-source software (https://gitlab.com/tjian-darzacq-lab/spot-on-matlab) based on a model previously introduced in (Mazza et al. 2012) and modfied in (Hansen et al. 2017) to exclude state transitions. In brief, Spot-On performs fits to the distribution of displacements at multiple frameshifts to a 2state kinetic model and provides estimates of the fraction of molecules bound and free, and the corresponding apparent diffusion coefficients for each state (Figure 3-figure supplement 1) and corrects for the probability of molecules diffusing out of the axial detection range. We performed fitting using the following parameters: Gaps Allowed: 1, Jumps to Consider: 4, TimePoints: 8, Observation Slice: 0.8 μm, Fit Iterations 5. The fit parameters for each data set are summarized in Figure 3-figure supplement 1. Data are represented as the mean over the 3 embryo replicates ± SEM.

### Calculation of residence times from long exposure single molecule trajectories

Imaging with sufficiently long exposure times effectively blurs out fast-moving molecules into the background while molecules stably bound for a significant duration of the exposure time are imaged as diffraction limited spots (Hansen et al. 2017; Watanabe and Mitchison 2002; Mir et al. 2017; Teves et al. 2016; J. Chen et al. 2014). Thus the trajectories from the 500 ms datasets are used to infer the genome average longlived (specific) binding times.

To infer the residence time, the length of trajectories in time is used to calculate a survival probability (SP) curve (1- cumulative distribution function of trajectory lengths). Since the SP curve contains contributions from non-specific interactions, slowly moving molecules, and localization errors a double-exponential function of the form SP(t)=F*(exp(-k_ns_*t))+(1-F)(exp(-k_s_*t))is fit to the SP curve, where k_ns_ is the off-rate for the short-lived (non-specific) interactions and k_s_ correspond to the off-rate of long lived (specific) interactions (J. Chen et al. 2014; Hansen et al. 2017; Mir et al. 2017) (Figure 3-figure supplement 2). For fitting probabilities below 10^-3 are not used to avoid fitting the data poor tails of the distribution. An objective threshold on the minimum number of frames a trajectory lasts is then used to further filter out tracking errors and slow-diffusing molecules (Mazza et al. 2012; Hansen et al. 2017). The objective threshold is determined by plotting the inferred slow rate constant and determining where values converge to a single value. Although the 500 ms Bcd data set converges at 2 frames (1 s), the Zld data set converges at 4 frames (2 s) (Figure 4-figure supplement 1B-C). The survival probability distribution for Zelda is likely dominated by shortlived interactions at shorter timescales and is most likely a reflection of the same complex mixed population (specific and non-specific DNA binding, along with another population whose motion is constrained perhaps by protein-protein interactions) we observed in the MSD curves (Figure 2B). Thus a 4 frame threshold was used for the calculation of the specific residence time.

Next, since the inferred k_s_ as described above is biased by photobleaching, and nuclear and chromatin movement, bias correction is performed using the His2B data as k_s,true_=k_s_-k_bias_, where k_bias_ is the slower rate from the double-exponent fit to the His2B SP curve as described previously (Teves et al. 2016; Hansen et al. 2017; J. Chen et al. 2014). This correction is based on the assumption that photobleaching, unbinding, and loss of trajectories from motion are all independent Poisson processes. The genome wide specific residence time is then calculated as 1/k_s,true_. The effectiveness of this bias correction is checked by calculating the residence time from both the 100 ms and 500 ms frame rate data and observing convergence to within 1 sec (Figure 4-figure supplement 1C).

### Fluorescence Recovery After Photobleaching

FRAP was performed on a Zeiss (Germany) LSM 800 scanning confocal microscope equipped with several laser lines, of which the 488 nm laser was used for all experiments described here. Images were collected using a Plan-Apochromat 63x 1.40 NA oil-immersion objective using a window 50.7 µm by 3.6 µm. Bleaching was controlled by the Zen software, and experiments consisted of 10 frames collected before the bleach and 1000 frames collected after at a frame rate of 24 ms. In each frame, five circular bleach spots of 1 µm diameter were chosen to be a sufficient distance from nuclear edges. The spots were bleached using maximum laser intensity, with dwell time adjusted to 0.57 µs, which was chosen because it gave a sufficiently deep bleach of Bicoid, the fastest-recovering molecule we studied. Total bleach time was 1.5 s.

We collected data from at least three embryos for each molecule studied. Nuclei in the early embryo are highly mobile, and we found that the most reliable method to find stable nuclei was to simply collect many movies and select the ones in which nuclei remain stable for the duration of the experiment. We collected movies with stable nuclei for a total of at least 50 bleach spots (50 nuclei) total for each molecule. To quantify and bleach-correct FRAP data, we used a custom-written MATLAB software pipeline. Briefly, for each frame we manually select several “dark” spots that are not within nuclei and several “control” spots that are within bleached nuclei but well-separated from the bleach spot. We use a 600 nm diameter circle to calculate the signal at the spots in order to make the measurement robust to small chromatin movements. For each frame, the mean of the dark spots was subtracted from the bleach spot values (background subtraction), and individual bleach spot values were divided by the mean of the control spots to correct for the reduction in total nuclear fluorescence. Finally, the values for each spot were normalized to its mean value for the ten pre-bleach frames. We observed that chromatin movement occasionally causes the bleach spot to drift far enough to affect the signal, so we manually curated resulting correct traces to remove anomalous spots. This culling resulted in 27-40 quality recovery curves for each molecule. These curves were averaged for each molecule, and the mean recovery curve was used in figures and fitting.

We fit resulting FRAP curves to the reaction-dominant model (Sprague et al. 2004):

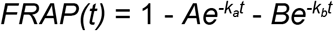

From these fits, we used the slower coefficient to estimate the time to half-recovery for the population of bound molecules.

### Analysis of single molecule binding in the context of Zld enrichment

To analyze single molecule trajectories of Bcd and Zld in the context of Zld enrichment first a relative enrichment map for each nucleus was calculated. For each reconstructed 100 ms exposure Zld image, first each nucleus was identified and segmented out of the image using an in house segmentation algorithm built in MATLAB (Figure 6-figure supplement 1). First the grayscale image was Gaussian filtered with a sigma of 5 pixels to enhance the contrast of the nuclei, the filtered image was then thresholded using the inbuilt adaptive threshold function in MATLAB with a sensitivity value set to 0.6. A morphological dilation was then performed on the binary mask using a disk structuring element with a radius of 3 pixels and multiplied with hand drawn mask to remove edges of the embryo and non-cortical regions deep where no nuclei were present or imaging contrast was low. Holes within the nuclei binary mask were then filled using the MATLAB imfill function. A label matrix was generated from the resulting binary mask and the size distributions and eccentricities of segmented regions were calculated, an area and eccentricity cutoff was then applied to remove false positives to generate the final label matrix. Label matrices were then further curated to remove false positives. A relative enrichment map was calculated for each nucleus individually by assigning each pixel in the nuclear value the percentile range it fell in over the entire distribution of intensity values in the nuclear area in with a resolution of 1 percentile. Each single molecule trajectory was then assigned a relative enrichment value based on the mean enrichment of the pixels it fell in during the course of the trajectory. From visual examination we determined that a 85 relative enrichment value threshold was reliable in differentiating the highest enriched Zld regions, corresponding to hubs, from the rest of the nucleus. Fold change in densities of detections were calculated by counting the total number of trajectories in areas of enrichment greater than 85 vs. the rest of the nucleoplasm. As the single molecule trajectories from this data set are limited in length to 1 s, an accurate estimate of the residence time from fits to the survival probability distribution could not be obtained as was done above.

### Analysis of protein distribution in context of transcription dynamics

Two-color 4D LLSM imaging was performed on embryos with the MS2-tagged *hb* BAC (Figure 7-figure supplement 1) and expresing MCP-mCherry crossed with either mNeonGreen-Zld or eGFP-Bcd embryos. Z-stacks of 61 slices were acquired with a spacing of 250 nanometers to cover a range of 15 µm with an exposure time of 80-100 ms in each channel at each slice. Images in both channels were acquired at each zposition sequentially before moving to the next slice. The time between each volume acquired was ~ 5 seconds, the total length of the acquisition varied but at least one complete nuclear cycle was imaged for each embryo. The field of view for each embryo was centered at between 25-35% of the embryonic length from the anterior tip of the embryo to ensure that all nuclei in the image were within the hunchback expression domain. Data from a total of 6 embryos each for mNeonGreen-Zld or eGFP-Bcd were analyzed.

To analyze the distribution of Zld or Bcd around sites of active *hb* transcription the signal from the MS2 site was used as a marker for the active locus. Each MS2 site was localized through a custom built detection software (Figure 7-figure supplement 1). First, the data was manually examined and annotated to simplify the segmentation procedure by only considering frames in which transcription was occuring. A 3D difference of gaussian image was then calculated at each frame to enhance the contrast of the MS2 site, a global threshold was then applied to generate a 3D binary mask for each frame. The binary mask was filtered to remove structures too big or too small to be from a MS2 site and a label matrix was generated. The xyz weighted center of each labelled region was then used to calculate line profiles extended 1 micron from the center of the region in each direction. The ratio of the maximum and minimum values in the line profile were used to determine if the labeled region was in a nucleus. This calculation is effective as in the MCP-mCherry channel the nucleoplasm around the MS2 site appears dark whereas in the remainder of the embryo the background is high, thus labeled regions with low contrast ratios were discarded. The maximum value of the profile of the remaining labelled regions was then used to localize the center of the active locus. The detected loci in each frame were then connected in time using a nearest neighbor algorithm.

The position list of the detected and tracked active loci were then used to crop a 2.18 µm window around the center of each locus in x-y, if any part of the window did not lie within a nucleus the locus was not considered for further analysis. For the remain loci a control window was cropped at a distance of 2.6 µm from the center of the locus in x-y. If a control window could not be found that did not completely lie within the nucleus the corresponding locus was also not considered for further analysis. In this manner a total of 3943 and 6307 windows centered around loci and corresponding control points were accumulated from the Bcd and Zld datasets respectively. The mean image at the locus was then calculated (Figure 7C and Videos 18 and 19) and a radial profile was calculated for Zld or Bcd centered at the active locus or the random control site. The radial profiles were then normalized to 1 at the maximum radius.

## Acknowledgments

We thank Robert Tjian for useful discussion and extensive advice on the science and the manuscript. We thank Jacques Bothma, Simón Álamos and Hernan Garcia for useful discussions and providing the *hb-MS2* and MCP-mCherry fly lines. We also acknowledge the advice and comments provided by all members of the Eisen, Darzacq, Tjian, and Garcia labs over the course of this work. This work was supported by a Howard Hughes Medical Institute investigator award to MBE; XD acknowledges support from California Institute of Regenerative Medicine (CIRM) grant LA1-08013 and the National Institutes of Health (NIH) grants UO1-EB021236 and U54-DK107980; MRS was supported by an American Cancer Society postdoctoral fellowship.

## Competing Interests

We have no competing interests to declare.

